# Combinatorial Regimens Augment Drug Monotherapy for SARS-CoV-2 Clearance in Mice

**DOI:** 10.1101/2023.05.31.543159

**Authors:** Irfan Ullah, Fanny Escudie, Ivan Scandale, Zoela Gilani, Gabrielle Gendron-Lepage, Fleur Gaudette, Charles Mowbray, Laurent Fraisse, Renée Bazin, Andrés Finzi, Walther Mothes, Priti Kumar, Eric Chatelain, Pradeep D. Uchil

## Abstract

Direct acting antivirals (DAAs) represent critical tools for combating SARS-CoV-2 variants of concern (VOCs) that evolve to escape spike-based immunity and future coronaviruses with pandemic potential. Here, we used bioluminescence imaging to evaluate therapeutic efficacy of DAAs that target SARS-CoV-2 RNA-dependent RNA polymerase (favipiravir, molnupiravir) or Main protease (nirmatrelvir) against Delta or Omicron VOCs in K18-hACE2 mice. Nirmatrelvir displayed the best efficacy followed by molnupiravir and favipiravir in suppressing viral loads in the lung. Unlike neutralizing antibody treatment, DAA monotherapy did not eliminate SARS-CoV-2 in mice. However, targeting two viral enzymes by combining molnupiravir with nirmatrelvir resulted in superior efficacy and virus clearance. Furthermore, combining molnupiravir with Caspase-1/4 inhibitor mitigated inflammation and lung pathology whereas combining molnupiravir with COVID-19 convalescent plasma yielded rapid virus clearance and 100% survival. Thus, our study provides insights into treatment efficacies of DAAs and other effective combinations to bolster COVID-19 therapeutic arsenal.

## Introduction

Severe Acute Respiratory Syndrome coronavirus-2 (SARS-CoV-2) is an enveloped positive-strand RNA virus of the *Coronaviridae* family that is the causative agent of the Coronavirus disease-19 (COVID-19) pandemic and has caused over 6.24 million deaths worldwide to date ^1–3^. Immunity provided by neutralizing antibodies (nAbs) elicited by several vaccines and treatment using nAb cocktails have significantly reduced the mortality rate caused by the COVID-19 pandemic ^1, 4–9^. However, SARS-CoV-2 variants of concern (VOCs; including Alpha, Beta, Gamma, Delta, Omicron), that mutate quickly to escape nAbs with enhanced transmissibility, have led to several waves of pandemic ^10,11^. We are experiencing accelerated evolution of virus, enhanced immune escape and transmission as well as a notable decrease in vaccine protection resulting in requirement for additional bivalent booster doses especially in the elderly population ^12,13^. Although long-term COVID and post-acute sequelae associated with SARS-CoV-2 are not negligible, no major variant of concern (VOC) has emerged in the past year. Furthermore, COVID-19 is no longer considered a global health emergency by WHO ^14^. While Omicron VOC and its XBB lineages (recombinant variants of Omicron BA.2.10.1 and BA.2.75). have caused milder disease and fewer acute deaths than previous VOCs, there is no guarantee that new variants will not emerge. It is important to note that Sarbecovirus subgenus consists of many zoonotic viruses that can potentially spread into human populations. Several members of this family have caused epidemics with much greater mortality rate, including SARS-CoV-1 (2002/3) and MERS (2012). The lessons learned from previous and current Coronavirus outbreaks demand that we continue to develop broad, highly efficient, and multi-pronged approaches to protect the human population from current and future outbreaks ^15^.

Several Direct acting antivirals (DAAs) repurposed from former antiviral programs have been approved or received emergency use authorization for COVID-19 treatment ^16^. They are essential to combating evolving VOCs, especially in the elderly, immunocompromised, and diseased population where vaccination fails to produce the desired protective response, as well as in vaccine-hesitant groups. Unlike nAbs, whose efficacy is compromised with emergence of VOCs, DAAs target conserved viral enzymes that play an essential role in virus replication and are expected to be overall effective despite the nAb-escape mutations in spike proteins of VOCs. However, equivalent efficacy cannot be guaranteed and must be ascertained through experimentation. Two classes of antiviral drugs that target viral RNA-dependent RNA polymerase (RdRp) (molnupiravir, favipiravir), and Main viral protease (nirmatrelvir) are currently available in the clinic and are approved by the FDA or are available for off-label use to treat high-risk adults with SARS-CoV-2. Molnupiravir (MK-4482, EIDD-2801, beta-D-N4-hydroxycytidine NHC), marketed by Merck and Ridgeback Biotherapeutics and developed at Emory University to treat influenza, is an orally available ribonucleoside analog ^17^. After oral administration, molnupiravir, isopropylester prodrug of N4-hydroxycytidine (NHC, EIDD-1931), is phosphorylated intracellularly to the active NHC triphosphate form. During virus replication, NHC triphosphate is incorporated into viral RNA by the viral RNA polymerase and then promotes misincorporation of guanosine or adenosine resulting in mutagenesis. The virus is ultimately rendered non-infectious and unable to replicate after accumulating deleterious errors throughout its genome which is also known as “error catastrophe”. Favipiravir (6-fluoro-3-hydroxypyrazine-2-carboxamine; available for off-label use) was originally discovered in a chemical screen for activities against influenza virus by Toyoma chemicals and has been approved in Japan for the management of emerging pandemic influenza infections since 2014 ^18, 19^. Favipiravir demonstrates broad spectrum antiviral activity against a variety of RNA viruses and is also a prodrug that is metabolized intracellularly into its active ribonucleoside 5′-triphosphate form that acts as a nucleotide analog to selectively inhibit RdRp and functions by inducing lethal mutagenesis or chain termination if two molecules of favipiravir are incorporated together. Several studies have reported *in vitro* inhibitory activity of favipiravir against SARS-CoV-2 ^19, 20^. Nirmatrelvir was originally developed by Pfizer in response to SARS-CoV-1 and targets the highly conserved Main protease/3C-like protease (Mpro, 3CL^PRO^) of SARS-CoV-1, SARS-CoV-2 and beta coronaviruses ^21, 22^. Mpro is involved in the processing of the viral polyprotein into functional proteins for the virus to form replication complexes. Nirmatrelvir-mediated inhibition of Mpro leads to the formation of non-functional viral proteins, preventing the virus from replicating and spreading. Several studies have shown that both molnupiravir and nirmatrelvir can effectively suppress acute virus replication and associated disease in the Syrian hamster, in ferrets and macaques infected with SARS-CoV-2 ancestral strain ^23–26^. Moreover, results from *in vitro* studies or in hamsters suggest that both drugs retain activity against multiple VOCs including Delta and Omicron variants ^27^. However, DAAs that act by inducing mutations can be a double-edged sword that promote accelerated virus evolution and eventual drug-resistance. Therefore, safe, and synergistic combinations of DAAs that ideally target different steps of virus replication may be required to effectively eradicate the virus.

As SARS-CoV-2 evolves, independent preclinical evidence for efficacy of antiviral drugs or their combinations *in vivo* are needed and must be generated due to accrued mutations, differences in replication rate, titers, and transmissibility of VOCs. In this regard, we have developed a non-invasive bioluminescence imaging (BLI) guided infection model to monitor virus replication and dissemination using SARS-CoV-2 NanoLuc luciferase (nLuc) reporters ^28–32^. A BLI-guided analyses can reduce the number of animals used in experiments and conform to the 3Rs (Refine, Reduce, Replace) of animal research. Furthermore, BLI allows monitoring the impact of antiviral regimens on virus replication and spread from lung to distal organs, such as the brain and GI tract whilst also capturing the ability of the tested interventions to distribute into the affected tissues which may differ due to pharmacokinetic differences and the activity of drug transporters such p-glycoprotein ^29^. In this study we applied our established BLI to visualize SARS-CoV-2 VOC (Delta and Omicron) spread and persistence for evaluating efficacy of therapeutic drugs in K18-hACE2 transgenic mice. These mice are highly susceptible to human-tropic SARS-CoV-2 variants, (except Omicron and its sublineages) and succumb to infection-induced mortality characterized by acute respiratory disorder syndrome (ARDS) due to infection in the lung followed by lethal encephalitis^33^. We evaluated the anti-SARS-CoV-2 activities of favipiravir, molnupiravir, nirmatrelvir or a combination of molnupiravir with nirmatrelvir, in combating the highly transmissible Delta and Omicron VOCs. While monotherapy regimens of favipiravir, molnupiravir or nirmatrelvir can reduce virus replication in lungs, combination therapy with molnupiravir and nirmatrelvir was significantly more efficacious in eliminating Delta and Omicron VOC-infected cells in our preclinical model analyses. The reduction and not elimination of virus infection under monotherapy might have consequences for virus evolution and emergence of drug-resistant VOCs. Inflammasome activation and pyroptosis-induced cell death mechanisms play a critical role in exacerbating COVID-19 ^34, 35^. Therefore, we also evaluated the synergistic potential of Caspase1/4 inhibitor (VX-765) in combination with short-term molnupiravir dosing under therapeutic settings. A combination of molnupiravir with VX-765 significantly improved survival as compared with monotherapy in addition to mitigating SARS-CoV-2-induced imbalanced inflammatory response and lung pathology. We also explored the potential of combining antiviral drugs with convalescent plasma (CCP) therapy that are relatively less expensive source of treatment option in many of Low- and middle- income countries (LMICs) where resources are very limited. Our analyses demonstrate that CCP with Fc-effector functions can effectively synergize with short-term molnupiravir treatment and rapidly clear established infection mice. Thus, our study provides valuable insights into treatment efficacy of approved drugs, and their combinations with inflammasome inhibitors or CCPs as part of the COVID-19 therapeutic arsenal.

## Results

### Comparative Efficacy of Favipiravir, Molnupiravir, Nirmatrelvir and the Combination of Molnupiravir with Nirmatrelvir Against Delta (B.1.617.2) VOC in K18-hACE2 Mice

We tested the therapeutic efficacy of off-label use/FDA-approved DAAs, favipiravir (600 mg/kg i.p.), molnupiravir (250 mg/kg; oral) or nirmatrelvir (650 mg/kg, oral) in K18-hACE2 mice challenged with reporter Delta VOC expressing NanoLuc luciferase (Delta-nLuc) (intranasal, i.n.; 1 x10^5^ FFU). Delta VOC is known to replicate to higher levels than other VOCs in the lungs ^36^. Pharmacokinetic analyses revealed steep drop in drug concentrations at 24 h in the serum after oral or intraperitoneal dosing **(Figure S1)**. Therefore, we administered the drugs 6 hpi followed by two doses per day (BID) until 6 dpi to effectively inhibit virus replication. Vehicle-treated mice were used as controls **(Figure 1A)**. Virus infection was monitored with BLI every two days and morbidity assessed using daily body weight measurements. BLI and subsequent quantification of nLuc flux (photons/sec) revealed that Delta-nLuc was able to spread from the nose to establish infection in the lungs of vehicle-treated mice by 2 dpi **(Figure 1B-D)**. The virus subsequently expanded in the lungs before spreading to the brain between 5-7 dpi, resulting in a concomitant loss in body weight loss and 100% mortality by 6-7 dpi **(Figure 1E-F)**. Analyses of inflammatory cytokine mRNA expression (*Il6, Ccl2, Cxcl10, Ifng, Il1b*) in the brain and lung revealed 10-1,000-fold induction in vehicle-treated mice compared to uninfected control **(Figure 1G-H)**. A single-dose treatment of a well-characterized nAb CV3-1 (12.5 mg/kg, i.p., 1 dpi) served as a positive control to compare the degree of virologic control engendered by the drug regimen. As expected from our previous studies ^29^, CV3-1 nAb treatment completely controlled Delta VOC replication as indicated by near-absence of nLuc signals in all tissues (non-invasively and after necropsy), and body-weight loss with 100% survival as well as baseline levels of inflammatory cytokines and nucleocapsid (N) mRNA expression.

**Figure 1.**
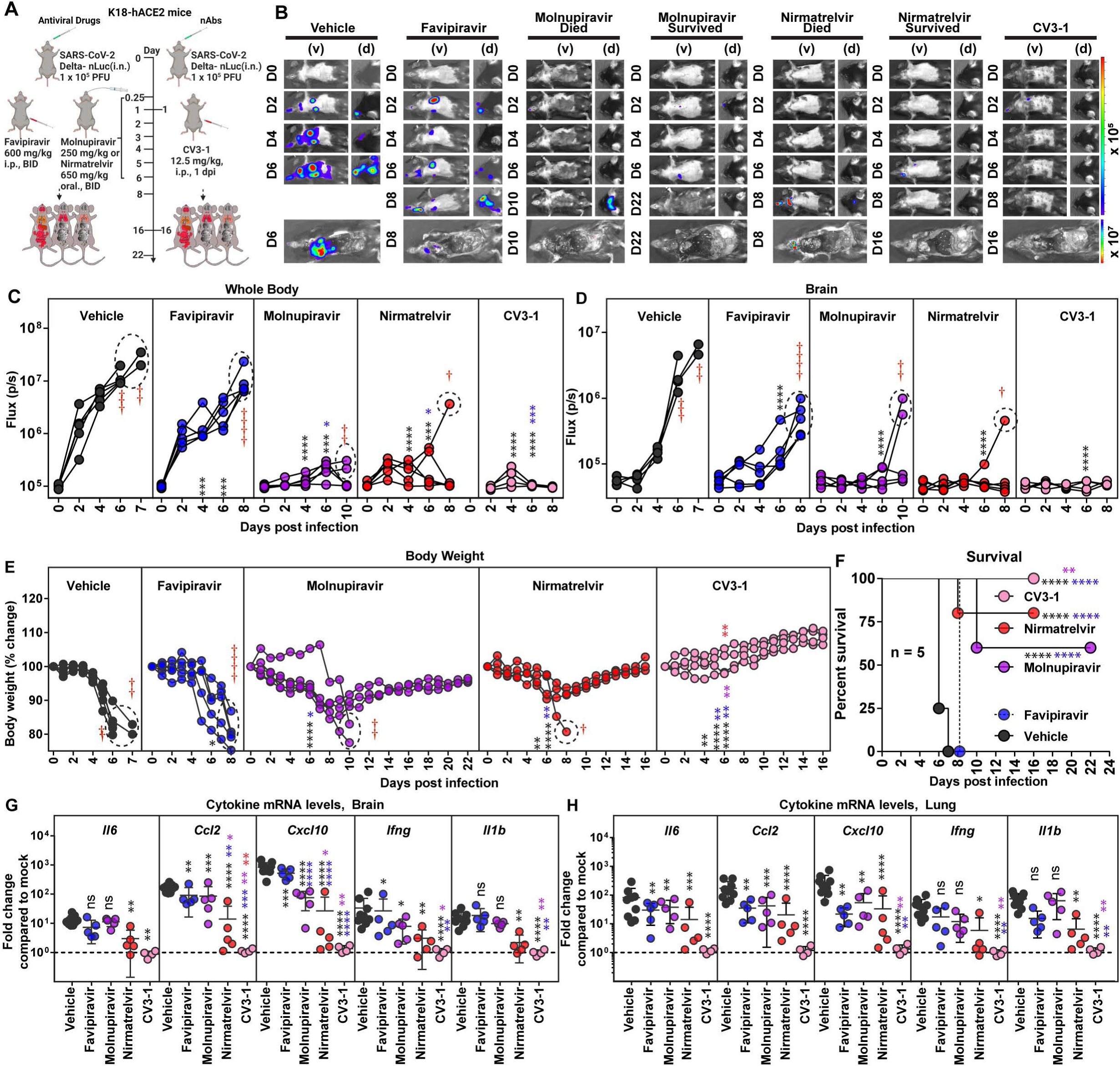
*In Vivo* efficacy of Favipiravir, Molnupiravir and Nirmatrelvir Monotherapy Regimen in K18-hACE2 Mice Against SARS-CoV-2 Delta VOC. (A) Experimental design for monitoring therapeutic efficacy of indicated drugs and neutralizing antibody (nAb) CV3-1 in K18-hACE2 mice challenged intranasally (i.n.) with 1 x 10^5^ PFU of reporter SARS-CoV-2-Delta-nLuc VOC. favipiravir (600 mg/kg body weight, b.w.) and nAb CV3-1 (12.5 mg/kg b.w.) were administered intraperitoneally (i.p.) while molnupiravir (250 mg/kg) and nirmatrelvir (650 mg/kg) given via oral gavage (oral) under a therapeutic regimen starting 0.25 dpi. All drugs except nAb (single dose) were given twice a day (BID) for 6 days. Vehicle-treated (Vehicle) mice (n=5-10) were used as control. (B) Representative BLI images of SARS-CoV-2-Delta-nLuc-infected mice under indicated treatment regimens in ventral (v) and dorsal (d) positions. Mice that succumb and survive challenge are shown separately for clarity. (C-D) Temporal quantification of nLuc signal as flux (photons/sec) computed non-invasively. (E) Temporal changes in mouse body weight with initial body weight set to 100% for an experiment shown in A. (F) Kaplan-Meier survival curves of mice (n = 5 per group) statistically compared by log-rank (Mantel-Cox) test for experiment as in A. (G-H) Fold changes in indicated cytokine mRNA expression in brain and lung tissues of mice under specified treatment regimens after necropsy upon death or at 14 or 22 dpi in the surviving animals. Data were normalized to *Gapdh* mRNA in the same sample and that in non-infected mice after necropsy. Grouped data in (C-E) and (G-H) were analyzed by 2-way ANOVA followed by Tukey’s multiple comparison tests. Statistical significance for group comparisons to Vehicle are shown in black, with favipiravir are shown in blue, with molnupiravir are shown in purple, with nirmatrelvir are shown as red and with CV3-1 are shown as pink. ∗, p < 0.05; ∗∗, p < 0.01; ∗∗∗, p < 0.001; ∗∗∗∗, p < 0.0001; ns not significant; Mean values ± SD are depicted. Points enclosed in dotted circles and red cross signs are used to indicate mice that succumbed to infection. See also Figure S2.

Although favipiravir therapy significantly reduced virus replication in the lungs (N mRNA expression, viral loads) and gut (nLuc flux), the inhibition was not enough to curb body-weight loss and eventual delayed spread of virus to the brain and nose as indicated by post-necropsy analyses **(Figure 1A-E, S2A-F)**. Consequently, all the mice in the cohort succumbed to Delta VOC-induced mortality with a one-day delay in death **(Figure 1F)**. Despite failing to achieve 100% survival, favipiravir treatment significantly reduced the mRNA expression of specific inflammatory cytokines (*Ccl2, Cxcl10*, 3-31-fold) in brain and (*Il6, Ccl2, Cxcl10*, 2-15 -fold) in lung tissues **(Figure 1G-H, Table 1)**. Comparatively, molnupiravir or nirmatrelvir monotherapy significantly reduced Delta VOC-induced morbidity (body-weight loss) and virus replication in target tissues (lungs, nose, brain, and gut) of surviving mice resulting in 60% and 75% survival, respectively, in addition to delaying death by 1-4 days in mice that succumbed to infection. BLI revealed that death of the mice in molnupiravir and nirmatrelvir-treated cohort was likely due to virus neuroinvasion **(Figure 1E-F, S2A-D)**. Post-necropsy analyses for tissue viral loads (nLuc flux, N mRNA expression, viral titers) in the nose, lung, brain, and gut also showed that molnupiravir or nirmatrelvir monotherapy significantly reduced tissue viral loads **(Figure S2A-D)**. Accordingly, we also observed significant reductions in all inflammatory cytokine mRNA expression in target tissues of molnupiravir or nirmatrelvir treated surviving mice **(Figure 1G-H)**. There have been reports documenting persistence of SARS-CoV-2 infection in the gut after respiratory clearance in pediatric and adult patients ^37–39^. However, as expected for orally delivered DAA, molnupiravir or nirmatrelvir-treated mice showed near baseline levels of nLuc signals especially in surviving mice compared to mice that succumbed to infection in all cohorts where we observed high level of SARS-CoV-2 infection in gut tissues **(Figure S2E, F)**. Notably, our analyses revealed that virus load in tissue homogenates on an average from mice under molnupiravir or nirmatrelvir monotherapy estimated on Vero E6 target cells (nLuc activity) were ∼92,000 and ∼4000-fold above baseline values respectively. Hence, although DAA monotherapy provided partial protection against Delta VOC, they did not achieve virus clearance to the same extent as nAb CV3-1 which displayed near baseline values (1.13-fold above background). Therefore, our analyses indicated a superior *in vivo* efficacy for nAb therapy compared to drug monotherapy regimens **(Table 1)**.

**Table 1.**
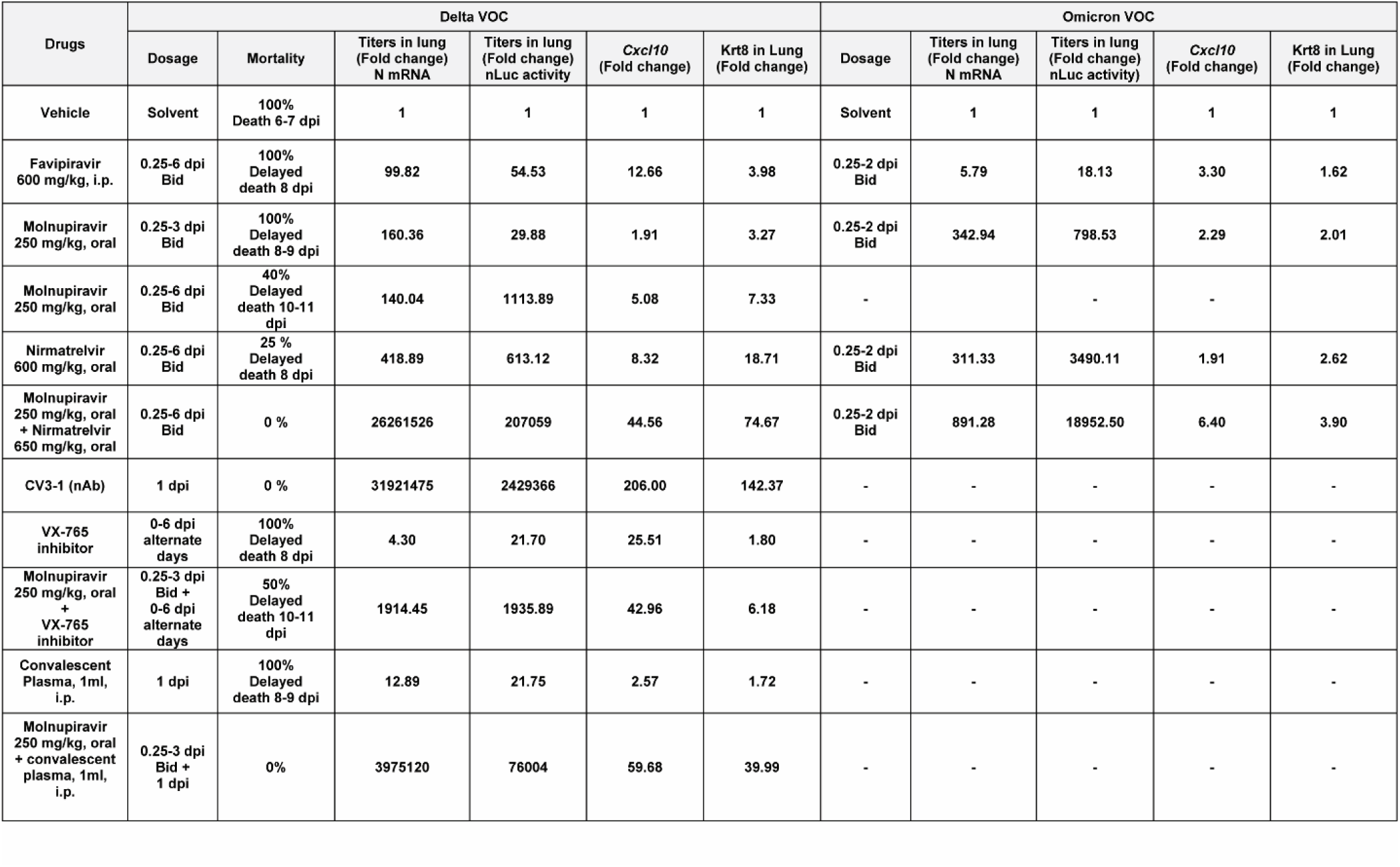
Summary of In Vivo Efficacy Analyses for Antiviral Therapeutic Regimens Evaluated in K18-hACE2 Mice Against SARS-CoV-2 Delta and Omicron VOC.

We next investigated if combination of DAAs, molnupiravir [250 mg/kg] and nirmatrelvir [650 mg/kg] (BID, for 6 days, starting 6 hpi) that target two distinct SARS-CoV-2 enzymes, polymerase, and the protease respectively, would demonstrate enhanced *in vivo* efficacy against Delta VOC **(Figure 2A)**. Strikingly and in contrast to vehicle-treated K18-hACE2 mice where virus replicated uncontrollably, the combination treatment significantly reduced virus replication below detection limits of BLI and prevented neuroinvasion suggesting successful elimination of virus-infected cells **(Figure 2B-D)**. These results were corroborated by prevention of body weight loss phenotype and morbidity resulting in 100% survival in mice receiving combination drug therapy versus vehicle-treated cohorts **(Figure 2E-F, Table 1)**. Moreover, post-necropsy analyses of viral loads in nose, lung, brain, and gut also indicated virologic control (N mRNA expression, nLuc activity) as well as significant reduction (5-450-fold) in inflammatory cytokine mRNA expression **(Figure 2G-I, S3A-E, Table 1)**. To assess lung pathology, we used *Krt8* mRNA expression, a marker for persistence of danger associated transitional progenitor (DATP) cells derived from alveolar epithelial cells during healing after lung injury ^40–42^. While monotherapy reduced lung pathology as measured by *Krt8* expression especially in surviving mice, CV3-1 nAb, and combination therapy were significantly superior in reducing lung injury **(Figure S4A-B)**. Thus, our results demonstrate that simultaneous inhibition of viral protease and polymerase activity is effective in clearing established pool of SARS-CoV-2-infected cells.

**Figure 2.**
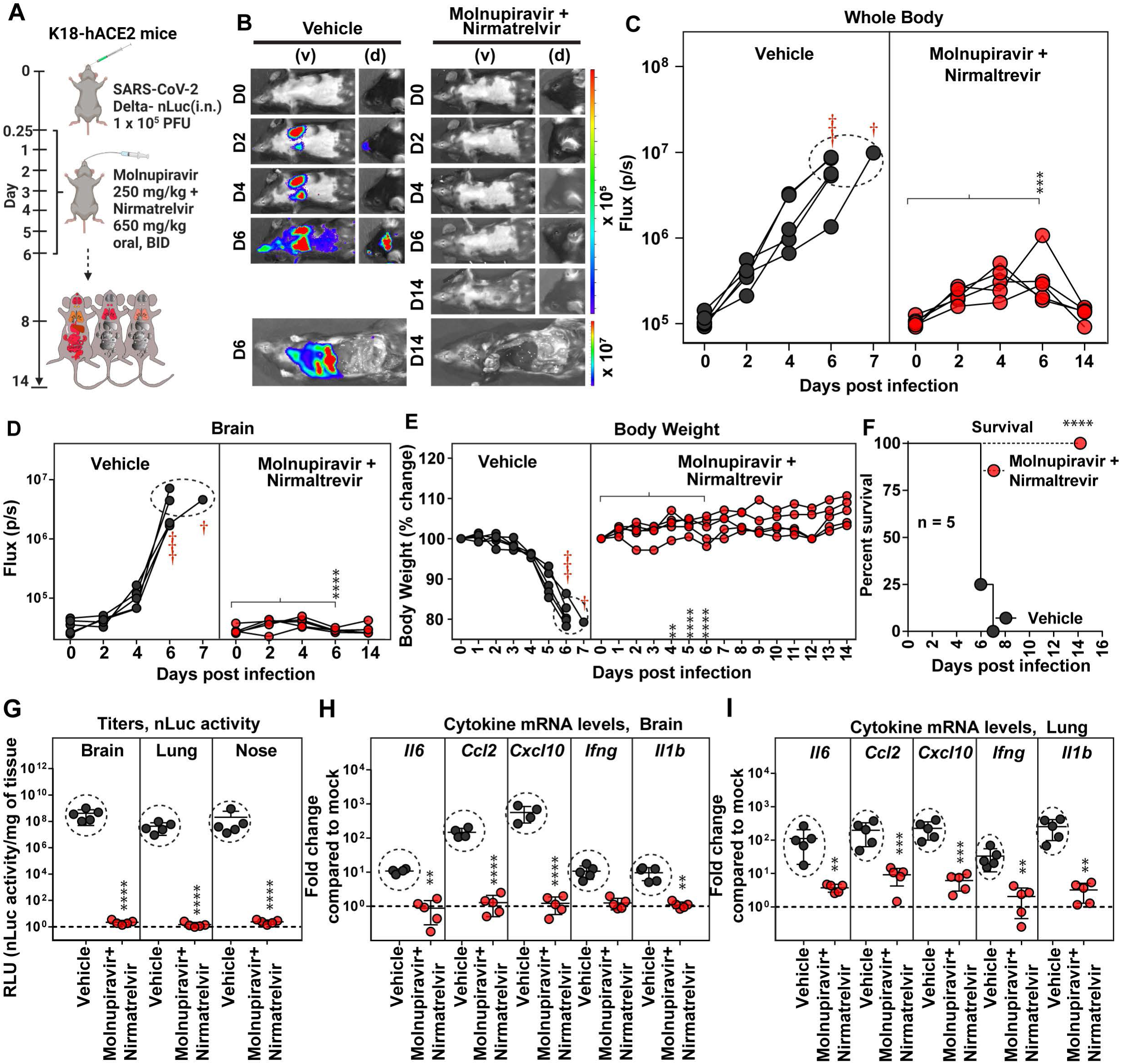
Treatment Efficacy of Molnupiravir and Nirmatrelvir Combination in K18-hACE2 Mice Lethally Challenged with SARS-CoV-2 Delta VOC. (A) Experimental design to test therapeutic efficacy of molnupiravir (250 mg/kg b.w.) with nirmatrelvir (600 mg/kg b.w.) in K18-hACE2 mice challenged with SARS-CoV-2 Delta-nLuc (1×10^5^ PFU, i.n.). The drugs were administered twice daily (BID) for 6 days beginning at 0.25 hpi. Vehicle-treated (Vehicle) mice (n=5) were used as control. (B) Representative BLI images of SARS-CoV-2-Delta-nLuc-infected mice in ventral (v) and dorsal (d) positions. (C-D) Temporal quantification of nLuc signal as flux (photons/sec) computed non-invasively. (E) Temporal changes in mouse body weight with initial body weight set to 100% for an experiment shown in A. (F) Kaplan-Meier survival curves of mice (n = 5 per group) statistically compared by log-rank (Mantel-Cox) test for experiment as in A. (G) Viral loads (nLuc activity/mg) in indicated organs from mice under specified treatment regimens evaluated using Vero E6 cells as targets when mice died from infection or at 14 dpi for surviving mice. (H-I) Fold changes in indicated cytokine mRNA expression in brain and lung tissues of mice under specified treatment regimens after necropsy upon death or at 14 dpi in the surviving animals. Data were normalized to *Gapdh* mRNA in the same sample and that in non-infected mice after necropsy. Grouped data in (C-E), and (G-I) were analyzed by 2-way ANOVA followed by Tukey’s multiple comparison tests. ∗, p < 0.05; ∗∗, p < 0.01; ns not significant; Mean values ± SD are depicted. Red cross signs and dots within indicated circles denote the mice that have succumbed to infection at specified times post infection. See also Figure S3.

### Comparative Efficacy of Favipiravir, Molnupiravir, Nirmatrelvir and the Combination of Molnupiravir with Nirmatrelvir Against Omicron (B.1.1.529) VOC in K18-hACE2 Mice

While Omicron VOC demonstrates high transmissibility, replication is attenuated in lungs of K18-hCE2 mice due to inefficient usage of TMPRSS2 ^43^. In addition, Omicron VOC infection is limited to nose, trachea and lungs and does not spread to brain in K18-hACE2 mice. This is manifested by non-lethal infection and only a transient loss in body weight with recovery after 3 dpi. *In vivo* efficacy studies of DAAs against Omicron VOC were therefore terminated at 3 dpi by treating mice at the same dosage as for Delta VOC challenges but from 0.25 to 2 dpi (BID) **(Figure 3A)**. We were able to monitor replication of Omicron-nLuc VOC after necropsy using whole body BLI and compared the effect of drug treatment on viral loads in nose, trachea, and lungs. As was the case with Delta VOC, monotherapy regimens were able to significantly reduce nLuc signals (2-10-fold) in the lung but did not eliminate virus-infected cells **(Figure 3B, C, Table 1)**. Among the three drugs evaluated, favipiravir was less efficient than molnupiravir or nirmatrelvir in preventing body weight loss and reducing virus loads in lungs (N mRNA expression and nLuc activity) compared to vehicle-treated cohorts of animals **(Figure 3D-H, Table 1)**. Notably, the mRNA expression of *Ifng* and *Il1b* in the lung and N mRNA in nose of all DAA-treated cohorts was significantly lower than vehicle-treated mice **(Figure 3I)**. Our data suggest that monotherapy with DAAs is effective against Omicron VOC in significantly reducing morbidity.

**Figure 3.**
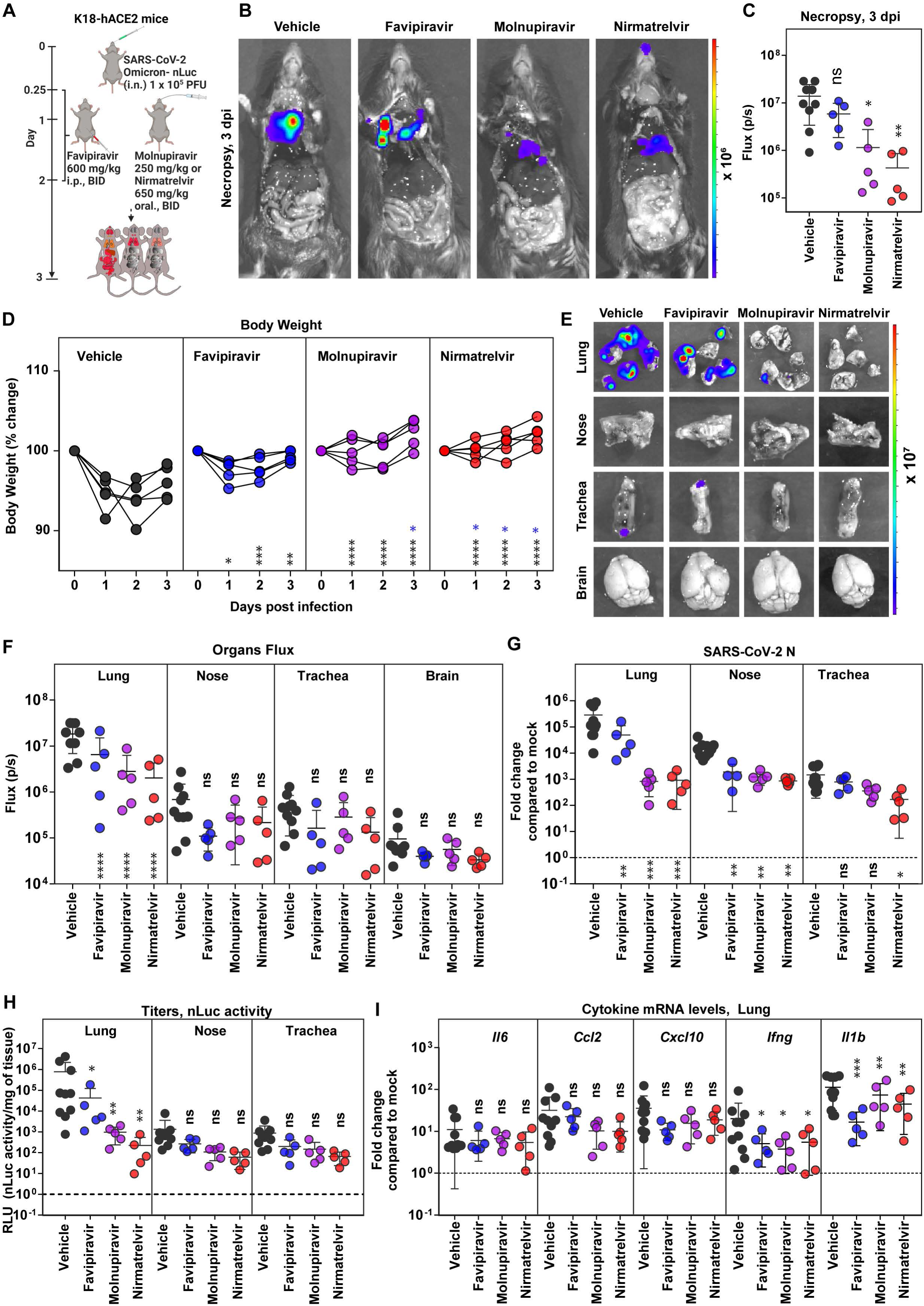
Efficacy of Favipiravir, Molnupiravir and Nirmatrelvir Monotherapy in K18-hACE2 Mice Against SARS-CoV-2 Omicron VOC. (A) Experimental design to test therapeutic efficacy of favipiravir (600 mg/kg b.w., i.p.), molnupiravir (250 mg/kg b.w. oral) and nirmatrelvir (650 mg/kg b.w., oral) against SARS-CoV-2-Omicron-nLuc (1 x 10^5^ PFU, i.n.) in K18-hACE2 mice. The drugs were administered twice daily (BID) starting 6 hpi (0.25 dpi) for 2 days. Vehicle-treated (Vehicle) mice (n=5-10) mice were used as control. (B) Representative BLI images of SARS-CoV-2-Omicron-nLuc-infected mice after necropsy (3 dpi). (C) Quantification of nLuc signal as flux (photons/sec) computed after necropsy at 3 dpi. (D) Temporal changes in mouse body weight with initial body weight set to 100% for an experiment shown in A. (E-F) *Ex vivo* imaging of indicated organs and quantification of nLuc signal as flux (photons/sec) after necropsy for an experiment as in A. (G) Fold changes in Nucleocapsid (N) mRNA expression in lung, nasal cavity, and trachea after necropsy at 3 dpi. Data were normalized to *Gapdh* mRNA in the same sample and that in non-infected mice. (H) Viral loads (nLuc activity/mg) from indicated tissues at 3 dpi using Vero E6 cells as targets. (I) Fold changes in indicated cytokine mRNA expression in lung tissues after normalization with *Gapdh* mRNA in the same sample and that in non-infected mice after necropsy. The data in (C) was analyzed by one-way ANOVA followed by Kruskal-Wallis’s test and grouped data in (D), (F-I) were analyzed by 2-way ANOVA followed by Tukey’s multiple comparison tests. Statistical significance for group comparisons to Vehicle are shown in black, with favipiravir are shown in blue, with molnupiravir are shown in purple, with nirmatrelvir are shown as red. ∗, p < 0.05; ∗∗, p < 0.01; ns not significant; Mean values ± SD are depicted.

We next tested if the combination therapy of molnupiravir [250 mg/kg] and nirmatrelvir [650 mg/kg] (BID, for 2 days, starting 6 hpi) can augment the antiviral effect against Omicron VOC like Delta VOC **(Figure 4A)**. Indeed, the combination therapy was highly effective in controlling Omicron VOC replication as ascertained by absence of body-weight loss phenotype as well as nLuc signals (BLI) after necropsy **(Figure 4B-D)**. Post-necropsy imaging of organs (lung, nose, trachea, and brain), viral load analyses (BLI, N mRNA and titers) and inflammatory cytokine mRNA expression (lung; *Cxcl10, Ifng* and *Il1b*) also revealed significant reduction in viral loads (70-18285-fold) and inflammation (2-6 -fold) compared to vehicle-treated cohorts **(Figure 4E-I, Table 1)**. This was corroborated by significantly reduced *Krt8* mRNA expression suggesting reduced lung pathology upon treatment with combination of drugs compared to vehicle and monotherapy regimens **(Figure S4C)**. Thus, the combination of molnupiravir and nirmatrelvir was more effective in controlling replication and spread of the highly immune evasive Omicron VOC than the DAA monotherapy evaluated in the study.

**Figure 4.**
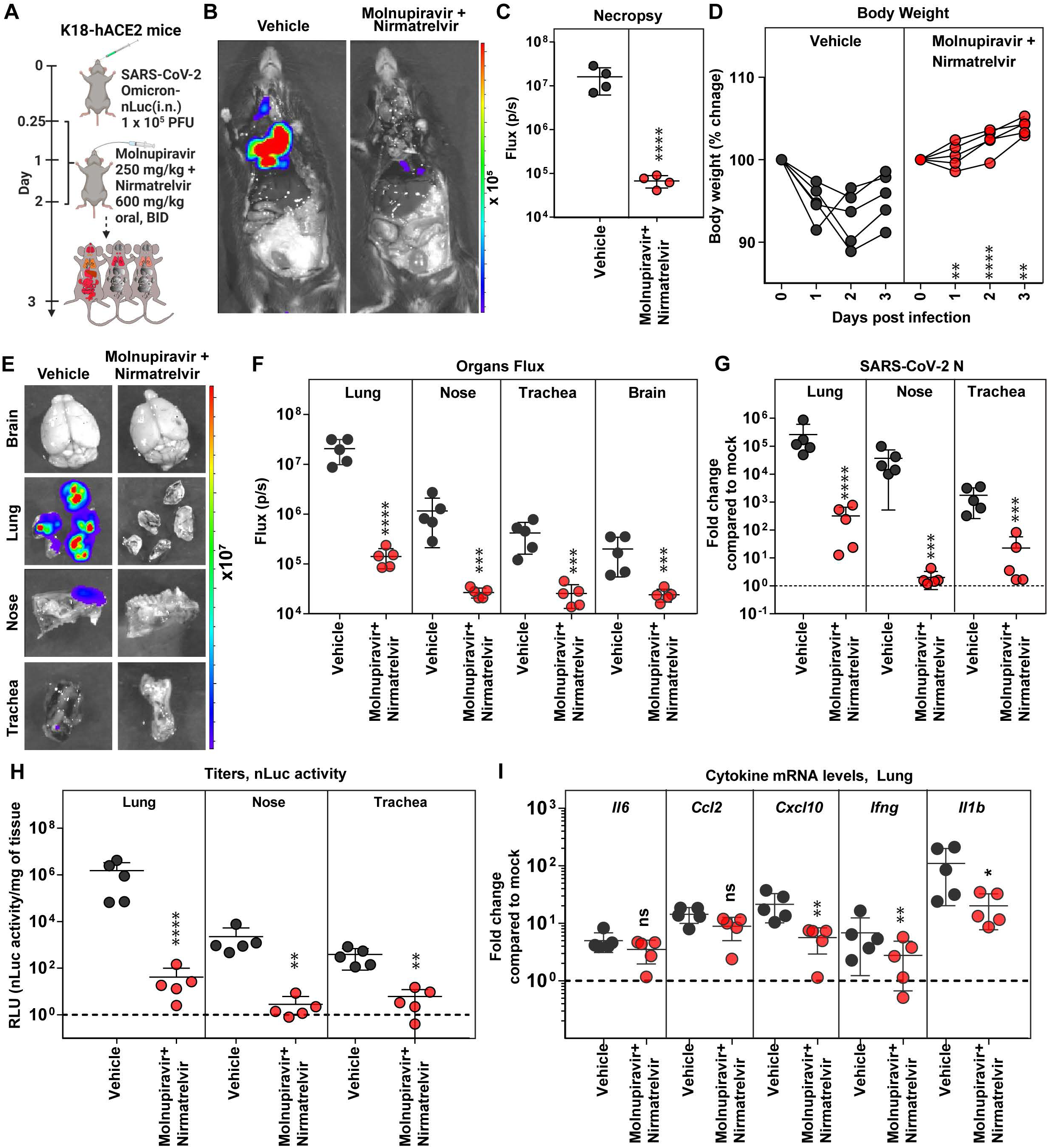
Therapeutic Efficacy of Molnupiravir with Nirmatrelvir in K18-hACE2 mice Against SARS-CoV-2 Omicron VOC. (A) Experimental design for testing efficacy of molnupiravir (250 mg/kg b.w., oral) with nirmatrelvir (600 mg/kg, b.w. oral) in K18-hACE2 mice challenged with SARS-CoV-2 Omicron-nLuc (1 x 10^5^ PFU; i.n.). The drugs were administered twice daily (BID) starting 6 hpi (0.25 dpi) for 2 days. Vehicle-treated (Vehicle) mice (n=5) were used as control. (B-C) Representative BLI images and temporal quantification of nLuc signal as flux (photons/sec) after necropsy at 3 dpi. (D) Temporal changes in mouse body weight with initial body weight set to 100% for an experiment shown in A. (E-F) *Ex vivo* imaging of indicated organs and quantification of nLuc signal as flux (photons/sec) after necropsy for an experiment shown in A. (G) Fold changes in nucleocapsid (N) mRNA expression in lung, nasal cavity and trachea tissues and treatment regimens at 3 dpi. Data were normalized to *Gapdh* mRNA in the same sample and that in non-infected mice after necropsy. (H) Estimated viral loads (nLuc activity/mg) from indicated tissues and treatment regimens using Vero E6 cells as targets at 3 dpi after necropsy (I) Fold changes in indicated cytokine mRNA expression for specified treatment regimens in the lung. Data were normalized to *Gapdh* mRNA in the same sample and that in non-infected mice. The data in (C) were analyzed by t test followed by non-parametric Mann-Whitney U tests. Grouped data in (D), (F-I) were analyzed by 2-way ANOVA followed by Tukey’s multiple comparison tests. ∗, p < 0.05; ∗∗, p < 0.01; ns; not significant. Mean values ± SD are depicted.

### Caspase 1/4 Inhibitor Improves Molnupiravir Therapeutic Efficacy and Extends Survival Against Delta VOC Challenge

SARS-CoV-2 infection elicits an imbalanced hyperinflammatory response primarily through activation of NLRP3-driven inflammasome pathway ^44, 45^. Mitigating SARS-CoV-2-induced inflammation alleviates lung pathology, whereas inhibiting inflammasome pathway reverses chronic lung pathology ^46–48^. We therefore explored if host-directed agents like VX-765 that specifically inhibit executor inflammatory Caspases 1 and 4 (i.p. 8 mg/kg, 0.25 - 6 dpi) which control cell death activated by inflammasomes ^49, 50^, can synergize with molnupiravir treatment regimen. To assess synergy, we chose a short-term 3-day molnupiravir treatment regimen (250 mg/kg, oral BID, 0.25-3 dpi) which was expected to be less efficient in inhibiting Delta VOC replication based on above 6-day administration experiments **(Figure 5A)**. Indeed, temporal BLI combined with nLuc signal quantification and body weight measurements revealed that although 3-day molnupiravir treatment significantly reduced Delta VOC replication in the lung, it only delayed neuroinvasion and subsequent death **(Figure 5B-D)**. Interestingly, we also observed reduced virus replication in the lung, delayed neuroinvasion (nLuc flux quantitation; 6 dpi) and a slower weight loss rate (6 dpi) in VX-765-treated mice that is not expected to directly inhibit virus replication compared to vehicle-treated cohorts **(Figure 5E)**. These data suggest that reducing inflammation can also reduce SARS-CoV-2 replication and spread **(Figure S5A-D)**. Despite the reduction in inflammatory cytokine mRNA expression and virus replication, all the mice in the VX- 765-treated cohort succumbed to Delta VOC-induced death **(Figure 5F)**. Moreover, we did not observe any significant reduction in lung pathology as assessed by *Krt8* mRNA expression in VX- 765 or short-term molnupiravir treated mice **(Figure S4B)**. Consequently, we explored if VX-765 and molnupiravir can be combined for potential synergy against lethal Delta VOC challenge. BLI revealed that the 3-day molnupiravir dosing, when combined with VX-765 treatment led to virologic control in 50% of the mice and a 3–4-day delay in neuroinvasion in the rest **(Figure 5B-D)**. Accordingly, half of the mice receiving the combination treatment displayed significant delay in body weight loss before succumbing to infection by 10-11 dpi while the other half survived **(Figure 5E-F)**. Post-necropsy analyses (BLI, RNA analyses) corroborated the virologic control in 50 % of the surviving mice in addition to significant overall reduction in inflammatory cytokine as well as *Krt8* mRNA expression (lung injury) in the lung **(Figure 5G-H, S5A-F, S4B, Table 1)** compared to individual and vehicle-treated mice. Thus, inhibition of the inflammasome pathway has the potential to augment monotherapy drug regimens and can have significant impact on preserving lung function.

**Figure 5.**
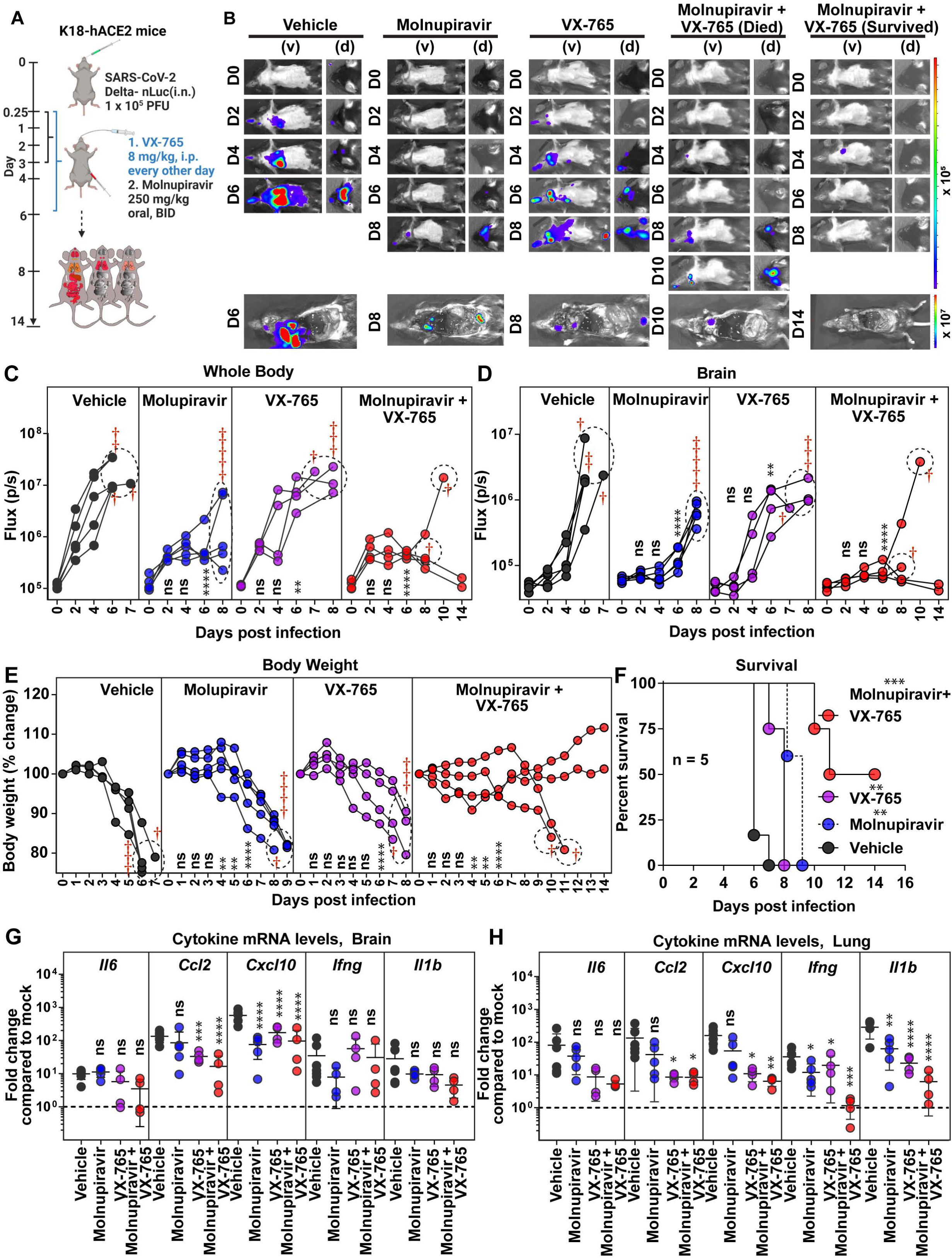
Caspase-1/4 Inhibitor can Augment Molnupiravir Therapeutic Efficacy in K18-hACE2 Mice Against Delta VOC. (A) Experimental design for testing efficacy of molnupiravir therapy (250 mg/kg b.w., oral) with Caspase 1/4 inhibitor (VX-765; 8 mg/kg b.w., i.p.) in K18-hACE2 mice challenged with SARS-CoV-2 Delta-nLuc (1 x 10^5^ PFU, i.n.). molnupiravir was administered twice daily (BID) starting 6 hpi (0.25 dpi) for 3 days and VX-765 was injected once every other day starting 6 hpi till 6 dpi Vehicle-treated (Vehicle) mice (n=5) mice were used as control. (B) Representative BLI images of SARS-CoV-2-Delta-nLuc-infected mice under indicated treatment regimens in ventral (v) and dorsal (d) positions. Mice that succumb and survive challenge are shown separately for clarity. (C-D) Temporal quantification of nLuc signal as flux (photons/sec) computed non-invasively. (E) Temporal changes in mouse body weight with initial body weight set to 100% for an experiment shown in A. (F) Kaplan-Meier survival curves of mice (n = 5 per group) statistically compared by log-rank (Mantel-Cox) test for experiment as in A. (G-H) Fold changes in indicated cytokine mRNA expression in lung and brain under specified drug regimens. RNAs were extracted when the mice succumbed to infection or for surviving mice at 14 dpi. Data were normalized to *Gapdh* mRNA expression in the same sample and that in non-infected mice after necropsy. Grouped data in (C-E), and data in (G-H) were analyzed by 2-way ANOVA followed by Tukey’s multiple comparison tests. Statistical significance for group comparisons to Vehicle are shown in black, with molnupiravir are shown in blue, with VX-765 are shown in purple, with molnupiravir+ VX-765 are shown as red. ∗, p < 0.05; ∗∗, p < 0.01; ns not significant; Mean values ± SD are depicted. Dotted circles with red cross signs at indicated time points are used to indicate mice that succumbed to infection. See also Figure S4.

### COVID-19 Convalescent Plasma (CCP) can Synergize with a Short-term Molnupiravir Treatment Regimen for Effective Virologic Control of Delta VOC

We next investigated if COVID-19 convalescent plasma (CCP) available in many countries, including those with limited resources (LMIC), can augment the short 3-day monotherapy with molnupiravir (250 mg/kg, oral BID, 0.25-3 dpi). We chose an ancestral strain elicited CCP (CCP-5, previously characterized in detail) ^32^ with robust Fc-effector activity required to eliminate persistent infection, but with low neutralizing activity against Delta VOC to mimic conditions resulting from immune evasion displayed by circulating VOCs. CCP-5 was administered only once at 1 dpi (i.p) either alone or in combination with molnupiravir **(Figure 6A)**. Longitudinal BLI and nLuc flux quantification revealed that although monotherapy using molnupiravir or CCP-5 reduced viral infection, delayed neuroinvasion and extended survival by 2-3 days, they failed to protect mice against Delta VOC-induced mortality **(Figure 6B-F)**. Notably, combining short-term molnupiravir treatment with CCP-5 resulted in rapid virologic control (nLuc flux quantification), significantly reduced morbidity as demonstrated by prevention of body weight loss and 100% survival **(Figure 6E-F)**. These data were corroborated by post necropsy analyses that showed elimination of nLuc signal as well as viral loads in all analyzed organs **(Figure S6A-F)**. We also observed a significant reduction in inflammatory cytokine mRNA expression in both the brain and the lung, as well as less lung damage (*Krt8* mRNA expression) in mice treated together with CCP-5 and molnupiravir **(Figure 6G-H, S4B, Table 1)**. Thus, combining a widely available resource such as CCP with a DAA can potentially improve antiviral drug therapies and achieve rapid virus clearance.

**Figure 6.**
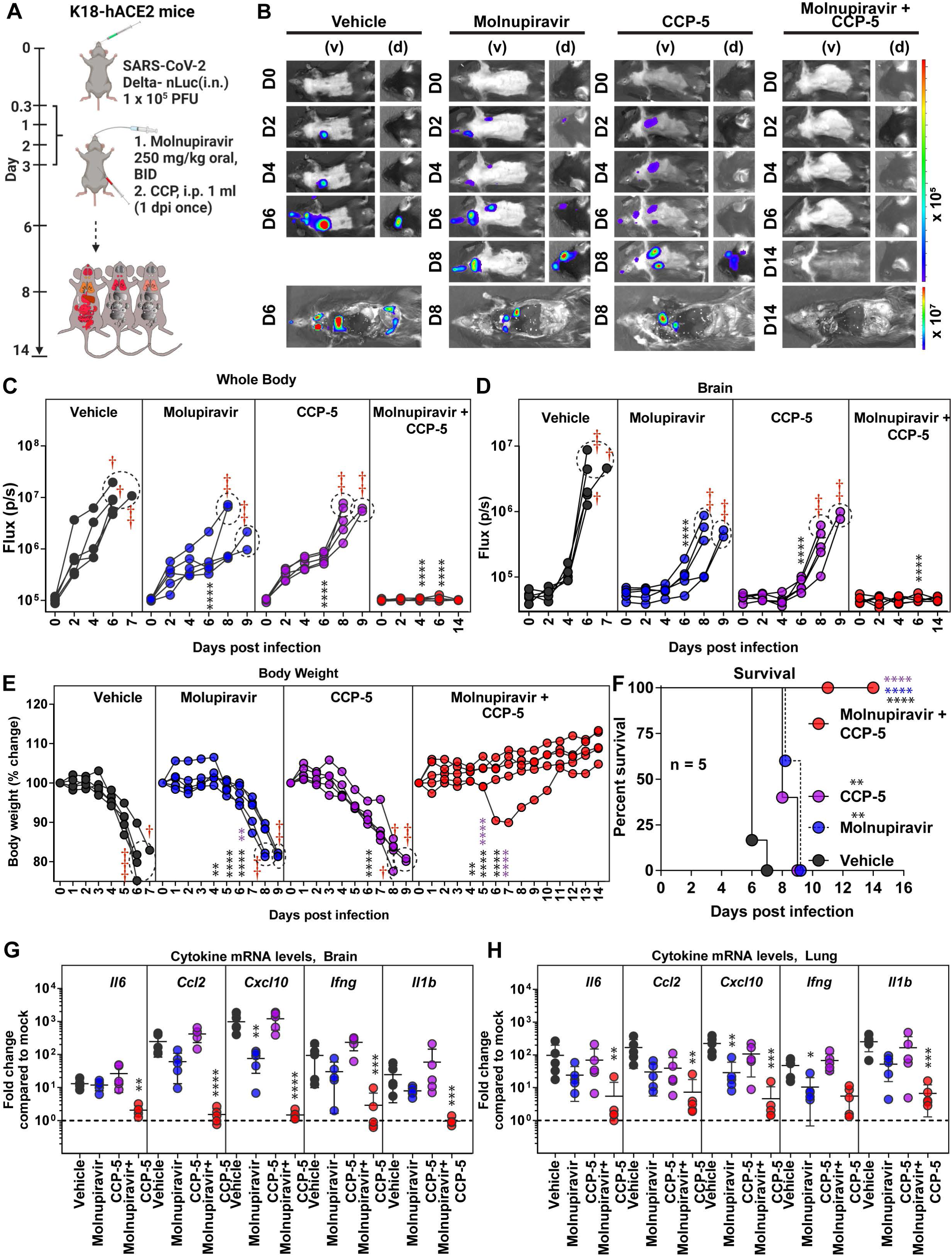
COVID-19 Convalescent Plasma 5 (CCP-5) can Augment Molnupiravir Therapeutic Efficacy in K18-hACE2 Mice Against Delta VOC. (A) Experimental design for testing synergy of molnupiravir therapy (250 mg/kg b.w., oral) with CCP-5 (i.p, 1ml, 1 dpi) in K18-hACE2 mice challenged with SARS-CoV-2 Delta-nLuc (i.n., 1 x 10^5^ PFU). Vehicle-treated (Vehicle) mice (n=5) mice were used as control. (B) Representative BLI images of SARS-CoV-2-Delta-nLuc-infected mice under indicated treatment regimens in ventral (v) and dorsal (d) positions. (C-D) Temporal quantification of nLuc signal as flux (photons/sec) computed non-invasively. (E) Temporal changes in mouse body weight with initial body weight set to 100% for an experiment shown in A. (F) Kaplan-Meier survival curves of mice (n = 5 per group) statistically compared by log-rank (Mantel-Cox) test for experiment as in A. (G-H) Fold changes in indicated cytokine mRNA expression in brain and lung tissues under specified treatment conditions. RNA was extracted from tissues of mice when they succumbed to infection or for surviving mice at 14 dpi. Data was normalized to *Gapdh* mRNA expression in the same sample and that in non-infected mice after necropsy. Grouped data in (C-E), and data in (G-H) were analyzed by 2-way ANOVA followed by Tukey’s multiple comparison tests. Statistical significance for group comparisons to Vehicle are shown in black, with molnupiravir are shown in blue, with CCP-5 are shown in purple, with molnupiravir combined with CCP-5 are shown as red. ∗, p < 0.05; ∗∗, p < 0.01; ns not significant; Mean values ± SD are depicted. Dotted circles with red cross signs at indicated time points are used to denote mice that succumbed to infection. See also Figure S5.

## Discussion

Antiviral interventions remain vital as SARS-CoV-2 VOCs may continue to emerge periodically with unpredictable clinical severity and ability to escape pre-existing spike-based vaccine immunity. It is therefore important to evaluate the antiviral drugs regimens and multi-targeted combination strategies that work independently from an effect on spike protein. DAAs, molnupiravir/favipiravir and nirmatrelvir which target SARS-CoV-2 polymerase and protease respectively are approved or available for off-label use as monotherapy for use in high-risk patients with COVID-19 ^51, 52^. These DAAs were discovered for use either against influenza or SARS-CoV-1 and were repurposed for use against SAR-CoV-2. Compared to molnupiravir, nirmatrelvir has superior in vivo efficacy (60% vs 75% survival) and is unlikely to induce host mutations as is the concern with molnupiravir. However, apart from improving the half-life through inhibition of Cytochrome P450 3A (CYP3A) family of oxidizing enzymes by combining with Ritonavir, a HIV protease inhibitor, accelerated improvement of nirmatrelvir subsided after SARS-CoV-1 outbreak ^53, 54^. While acknowledging that further development would have helped significantly in countering the current SARS-CoV-2 pandemic, this was expected due to lack of patients for human trials and available funding. In this regard, preclinical animal models of infection play a crucial role in developing promising antiviral candidates into near-finished products as well as identifying synergistic combinations for therapeutic use that are essential to pandemic preparedness.

Here we combined a stringent K18-hACE2 model of SARS-CoV-2 infection with BLI to track highly pathogenic Delta VOC and immune invasive Omicron VOC replication for studying efficacies of antivirals drugs as monotherapy and dual therapy. Our analyses revealed that nirmatrelvir had better *in vivo* efficacy than molnupiravir, while favipiravir was the least effective. Furthermore, molnupiravir monotherapy was more efficacious when administered over a six-day period than over a shorter three-day period (60% vs 0% survival). These data support a longer course of monotherapy to achieve virologic control. A longer duration may be especially important for treatment of patients with long-term COVID. While none of the DAAs under monotherapy could match the *in vivo* efficacy offered by the nAb CV3-1, the combination of molnupiravir with nirmatrelvir showed promise in achieving reduction in virus loads that came close to nAb therapy for the highly pathogenic Delta VOC. The combination also showed better efficacy against Omicron VOC than monotherapy regimens. The inability to eliminate the virus during monotherapy treatment may have consequences for virus evolution and emergence of drug-resistant VOCs especially for mutagenic DAAs like molnupiravir and favipiravir. Suboptimal concentrations have been shown to support emergence of SARS-CoV-2 mutants in hamster models ^55^. Furthermore, widespread use of molnupiravir has led to circulation of SARS-CoV-2 virus populations that carry molnupiravir-induced signature mutations (C to T and G to A) in the United Kingdom, the United States of America, and Australia ^56^. However, despite this concern, evidence that these random mutations induced by mutagenic DAAs contribute to evolution of SARS-CoV-2 variants with drug-resistance or enhanced fitness is lacking ^56, 57^. These findings highlight the importance of developing a panel of DAAs that can match the rapid virus clearance mediated by nAbs and, preferably, be used in combinations targeting distinct stages of the viral life cycle to remain ahead of the expected emergence of drug-resistance mutations as has been implemented for treatment of HIV with combinatorial antiretroviral therapy.

An exacerbated inflammatory response is the hallmark of SARS-CoV-2 infection and reducing inflammation through use of corticosteroids like dexamethasone was beneficial in patients with severe COVID-19 that required respiratory support ^58^. In this regard, we found that mitigating host inflammasome activation through use of Caspase-1/4 inhibitor (VX-765) augmented recovery and survival (0% vs 50%) of lethally challenged K18-hACE2 mice when combined with DAA like molnupiravir over individual treatment regimen. These data demonstrate the potential of synergistically combining DAAs with host-directed inhibition to augment virus clearance and recovery. Interestingly, reducing inflammation using VX-765 resulted in reduced viral loads suggesting that inflammation contributes to conditions beneficial for SARS-CoV-2 replication *in vivo*. Consequently, VX-765 treatment alone also reduced viral loads in addition to DAA treatment. These data suggest a model where individual DAA require help from a second virus load-reducing measure. This can be accomplished either via combination therapy with a second DAA or with host-directed treatments such as VX-765.

Importance of multipronged strategy was further reinforced by data from treatment with a combination of DAA and convalescent plasma. Notably, a nAb operates like a combination therapy. The neutralizing activity targets the virus directly like a DAA while the Fc effector function adds a second line of attack against both free virus particles (through complement, antibody dependent cellular phagocytosis), and virus-producing infected cells (antibody-dependent cellular cytotoxicity) through innate immune cell engagement. While efficacy to neutralize SARS-CoV-2 VOCs by Abs diminishes due to escape mutations, the polyclonal Fc-effector properties demonstrated resilience since spike-binding antibodies can continue to bind via conserved or non-neutralizing epitopes ^59–61^. Therefore, we also explored the possibility of combining short-term molnupiravir treatment with ancestral strain-elicited convalescent plasma-5 that has robust Fc-effector activity required to clear infected cell reservoir especially during therapeutic interventions^32^. Indeed, the combination with CCP-5 significantly enhanced efficacy of molnupiravir and resulted in virologic control, reduced morbidity, and inflammatory response. Thus, Fc effector functions provided through CCP offer a second mechanism to reduce viral loads in addition to DAA monotherapy. Conversely, the diminished neutralizing ability of polyclonal Abs in CCP-5 is complemented by direct acting molnupiravir. Thus, our study provides a general model for efficacies of currently approved drugs and suggest a requirement for combinatorial therapeutic regimen, either a second DAA, or combinations with inflammasome inhibitors or CCPs to be effective against COVID-19.

## Limitations of the Study

We used K18-hACE2 mice as our model for *in vivo* efficacy analyses as they are highly sensitive to SARS-CoV-2 infection. By setting a high threshold for evaluating *in vivo* efficacy, the model allows identification of antiviral interventions with better translational potential. However, K18-hACE2 mice succumb due to virus neuroinvasion. While virus neuroinvasion remains debatable in humans, a more recent study has demonstrated that SARS-CoV-2 can infect astrocytes in human brain ^62–64^. In addition, acute and long-term neurological dysfunction are one of the most frequent extrapulmonary complications in 30% of COVID-19 patients and there is strong evidence of brain-related abnormalities even in patients with mild COVID-19 disease ^65, 66^. Our study revealed the utility of a CCP to complement the action of molnupiravir to achieve virologic control in mice. However, CCPs are complex compared to purified nAbs and necessitate careful evaluation and additional clarity towards utility of CCP with drugs like molnupiravir can only be provided by a clinical trial. Alternatively, a combination of well-studied and safe spike-binding Ab with robust Fc-effector activity together with DAAs can also be tested to harness the rapid virus clearance mediated by Abs for minimizing lung pathology which remains a concern even after virus is cleared especially in the aged population.

## ACKNOWLEDGEMENTS

This project was funded through DNDi under the support by the Wellcome Trust Grant ref: 222489/Z/21/Z through the “COVID-19 Therapeutics Accelerator” and subcontracted to P.D.U, NIH grant R01AI163395 to W.M., and by a CIHR operating Pandemic and Health Emergencies Research grant #177958 to A.F. A.F. is the recipient of Canada Research Chair on Retroviral Entry no. RCHS0235 950-232424.

## AUTHOR CONTRIBUTIONS

Conceptualization, P.D.U., E. C. F. E, I. S, C.M., L.F; Methodology, P.D.U., I.U., F.E., I.S., E.C.; Investigation, I.U., Z G., P.D.U.; Writing – Original Draft, P.D.U., I.U.; Writing – Review & Editing, P.D.U., A.F., W.M., P.K., R.B., P.B., M.C., I.U., E.C:, F.E., I.S., C.M., L.F.; Funding Acquisition, E.C, C.M., L.F., P.D.U, W.M., A.F.; Resources, W.M., P.K., A.F., and R.B.; Supervision, P.D.U., E. C., L.F. C.M., P.K.

## Competing interest

The authors declare no competing interests

## STAR METHODS

### RESOURCE AVAILABILITY

#### Lead Contact

Requests for resources and reagents should be directed to and will be fulfilled by the Lead Contact, Pradeep Uchil.

#### Materials Availability

All unique reagents generated in this study are available from the lead contact with a completed Materials Transfer Agreement

#### Data and Code Availability

All data reported in this paper will be shared by the lead contact upon request. This paper does not report the original code.

Any additional information required to reanalyze the data reported in this paper is available from lead contact upon request.

### EXPERIMENT MODEL AND DETAILS

#### Cell and Viruses

Vero E6 (CRL-1586, American Type Culture Collection (ATCC), were cultured at 37°C in RPMI supplemented with 10% fetal bovine serum (FBS), 10 mM HEPES pH 7.3, 1 mM sodium pyruvate, 1× non-essential amino acids, and 100 U/ml of penicillin–streptomycin. SARS-CoV-2 B.1.617.2 (Delta) and Omicron VOC was isolated from a patient in Yale New Haven Hospital. All viruses were sequence confirmed using Next Genome Sequencing (Yale Keck Facility). We generated nanoluc luciferase (nLuc) expressing reporter viruses for Omicron and Delta VOC for non-invasive BLI imaging of infected mice using circular polymerization extension reaction (CPER) as described previously ^67^. Briefly, viral RNA was converted into cDNA using PrimeScript RT kit (Takara Bio) using a mix of random and oligo dT primers. The cDNA was then used as the template to amplify 11 overlapping fragments that were ∼2-3 kb long using specific primers and PrimeStar GXL polymerase (Takara Bio). A NLuc-P2A cassette was inserted at the N-terminus of ORF8 (nLuc-P2A-Orf8). The gel-purified overlapping fragments were circularized by CPER using a Hepatitis Delta virus ribozyme-spacer-CMV promoter cassette that overlapped with the 3’ and 5’end of the genome. CPER reactions were transfected into a T-25 flask of HEK293 cells using polyethyleneimine and cultured under BSL3 conditions. The following day the HEK293 cells were resuspended and co-cultured with Vero E6 ACE2/TMPRSS2 (Vero AT) cells for 5-9 days until CPE was clearly visible and nLuc activity was detected in the culture supernatants. Delta-nLuc and Omicron-nLuc was propagated in Vero AT cells by infecting them in T150 cm^2^ flasks at a MOI of 0.1. The culture supernatants were collected after 18-24 h when cytopathic effects were clearly visible. The cell debris was removed by centrifugation and filtered through 0.45-micron filter to generate virus stocks. Viruses were concentrated by adding one volume of cold (4 °C) 4x PEG-it Virus Precipitation Solution (40 % (w/v) PEG-8000 and 1.2 M NaCl; System Biosciences) to three volumes of virus-containing supernatant. The solution was mixed by inverting the tubes several times and then incubated at 4 °C overnight. The precipitated virus was harvested by centrifugation at 1,500 × g for 60 minutes at 4 °C. The concentrated virus was then resuspended in PBS then aliquoted for storage at −80°C. All work with infectious SARS-CoV-2 was performed in Institutional Biosafety Committee approved BSL3 and A-BSL3 facilities at Yale University School of Medicine.

#### Ethics statement

CCP-5 was obtained from individual who was infected during the first wave of the pandemic, after at least fourteen days of resolution of COVID-19 symptoms^68^. The participant consented to the study (Héma-Québec, CER #2020–004). Research adhered to the standards indicated by the Declaration of Helsinki. All participants were adults and provided informed written consent prior to enrollment in accordance with Institutional Review Board approval.

#### Plasma samples

Recovered COVID-19 patient who had received a COVID-19 diagnosis by the Québec Provincial Health Authority and met the donor selection criteria for plasma donation in use at Héma-Québec were recruited. The participant was allowed to donate plasma at least 14 days after complete resolution of COVID-19 symptoms. A volume of 500 mL to 750 mL of plasma was collected by plasmapheresis (TRIMA Accel, Terumo BCT). Disease severity (date of symptoms onset, end of symptoms, type, and intensity of symptoms, need for hospitalization/ICU) was documented using a questionnaire administered at the time of recruitment. For additional details of CCP-5 (sex, age, blood group of the convalescent donor and day of collection post infection, please refer to key resource table.

#### Mouse Experiments

All animals were maintained in the (SPF-free) barrier facility of the Yale University Animal Resource Centre (YARC) within a 14:10 light: dark cycle. Breeding population of mice and infected animals are maintained in separate rooms. All SARS-CoV-2-infected animals were housed in animal room under BSL3 containment. Cages, animal waste, bedding, and animal carcasses were disposed and decontaminated following the guidelines of Environmental Health Services at Yale. All replication competent virus-infected animals were handled under ABSL3 conditions with personnel’s donning pressurized air purified respirators (PAPR), double gloves, shoe covers, sleeve covers and disposable gowns. All experiments described here were approved by Institutional Animal Care and Use Committees (IACUC) as well as SOPs approved by Institutional Environmental Health and Biosafety committee. hACE2 transgenic B6 mice (heterozygous) were obtained from Jackson Laboratory. 6–8-week-old male and female mice were used for all the experiments. The heterozygous mice were crossed and genotyped to select heterozygous mice for experiments by using the primer sets recommended by Jackson Laboratory. Each cohort size was n = 4–8 to allow statistical testing and conducted as 2–3 biological replicates (n = 2–3 per replicate) to allow parallel evaluation of different cohorts. The number of animals (n = 4–8 per cohort) needed to achieve statistically significant results were calculated based on a priori power analysis. We calculated power and sample sizes required based on data from pilot experiments and previous studies ^29–32, 69^. Animals with sex-and age-matched littermates were included randomly in the experiments. No animals were excluded due to illness after the experiments. At the time of experimentation, care was taken to include equal numbers of male and female mice whenever possible to ensure that sex of the animals does not constitute a biological variable during analysis.

### METHOD DETAILS

#### SARS-CoV-2 infection and treatment conditions

For all *in vivo* experiments, 6 to 8 weeks male and female mice were intranasally challenged with 1 x 10^5^ PFU SARS-CoV-2-nLuc Delta or Omicron-nLuc VOCs in 25-30 µL volume under anesthesia (0.5 - 5 % isoflurane delivered using precision Dräger vaporizer with oxygen flow rate of 1 L/min). The starting body weight was set to 100%. For survival experiments, mice were monitored every 8-12 h starting six days after virus challenge. Lethargic and moribund mice or mice that had lost more than 20 % of their body weight were sacrificed and considered to have succumbed to infection for Kaplan-Meier survival plots. Mice were considered to have recovered if they gained back all the lost weight.

Favipiravir was prepared at concentration of 50 mg/ml in 0.3% (w/v) sodium bicarbonate solution made in water. The drug was dissolved for 1h at room temperature by shaking in a thermomixer. favipiravir solution was prepared fresh on the day of administration for dosing twice at 10-12 h intervals. favipiravir administration was initiated at 6 hpi via intraperitoneal injection (i.p., 26 gauge) (600 mg/kg body weight) followed by twice daily administrations (BID) at 1200 mg/kg body weight/day) at 10-12 h intervals till 6 dpi.

Molnupiravir was dissolved in solution (Vehicle) of 10% PEG400 and 2.5% Kolliphor-EL in sterile milli Q water by shaking in thermomixer for 15 minutes and then by vortexing for 5 minutes. The solution was kept in thermomixer till it was ready for dosing. molnupiravir solution was prepared fresh daily. molnupiravir (250 mg/kg body weight) was made in a volume of 150 µl and administered orally using an oral gavage needle (20 gauge) starting at 6 hpi. On the following days, 250 mg/kg body weight of molnupiravir was administered two times daily (BID, 10-12 hours apart) until 3 dpi or 6 dpi (total dose of 500 mg/kg per day).

Nirmatrelvir was prepared fresh daily by dissolving the drug in 0.5% (w/v) aqueous methyl cellulose in 2% Tween 80 prepared in MilliQ water. The drug was dissolved by shaking in thermomixer for 2 h at 37 °C. nirmatrelvir was dosed at 650 mg/kg body weight in a volume of 250 µl orally using an oral gavage needle (20 gauge) starting at 6 h post infection. On the following days, 650 mg/kg body weight of nirmatrelvir was administered two times a day (BID, total dose of 1300 mg/kg body weight per day) (10-12 h apart) till 6 dpi.

To test efficacy of nAb against Delta VOC, nAb CV3-1 was administered once (12.5 mg/kg, i.p.) at 1 dpi using a 26-gauge needle. Caspase-1/4 inhibitor (VX-765) (InvivoGen, *in vivo* grade) was injected intraperitoneally at 8 mg/kg body weight starting at 6 hpi followed by every other day till 6 dpi. VX-765 was administered either alone (VX-765 cohort) or in combination with molnupiravir (250 mg/kg body weight). 1 ml of COVID-19 convalescent plasma-5 (CCP-5) ^32^ was administered intraperitoneally once at 1 dpi either alone (CCP-5 cohort) or in combination with molnupiravir (250 mg/kg body weight, oral). molnupiravir was administered till 3 dpi as mentioned above for all experiments designed to test synergy with VX-765 or CCP-5.

#### Pharmacokinetics of antiviral drugs

favipiravir (600 mg/kg body weight, i.p) and nirmatrelvir (650 mg/kg body weight, oral) was administered in K18-hACE2 transgenic as described above. The whole blood samples (150µl per time point) were collected at 0, 4 and 24 h post administration. The samples were then allowed to clot by leaving undisturbed at room temperature for 4 h. Neat clear serum was collected by sedimenting the samples at 14,000 x g for 20 min at room temperature. At the 24 h timepoint mice were sacrificed and tissue samples (brain, nose, lung, trachea, and gut) were collected after necropsy. Tissue samples were accurately weighed and were homogenized in 800 µL of 1x Dulbecco’s PBS in 2 mL tube containing 1.5 mm Zirconium beads in a BeadBug 6 homogenizer (Benchmark Scientific, TEquipment Inc). The homogenized tissues were then sedimented at 13,000 rpm for 20 min at 4°C and the clear supernatant was frozen and stored at −80°C until further analysis.

#### Mass spectrometry

Favipiravir was extracted from mouse serum and tissue homogenates using protein precipitation as sample preparation technique. 200 µL of internal standard (IS) solution (500 ng/mL of ^13^C_5_-ribavirin in acetonitrile) was added to an aliquot of 25 µL of sample (serum or tissue homogenate). The sample was vortexed for approximately 5 seconds and let stand for a period of 10 minutes, then centrifuged at 16,000 x *g* for 10 minutes and the supernatant was transferred to an injection vial for analysis The analysis was performed using a Thermo Scientific TSQ Quantiva triple quadrupole mass spectrometer interfaced with a Thermo Scientific Ultimate 3000XRS UHPLC system using a heated electrospray ion source (HESI). MS detection was performed in negative ion mode, using selected reaction monitoring (SRM). The precursor-ion reactions for favipiravir and IS were set to 156.1 → 113.1 and 248.1 → 111.1, respectively. Isocratic elution was used with an Agilent Poroshell HILIC-Z column (100*2.1 mm I.D., 2.7µm) and guard cartridge operating operating at 45°C. The mobile phase conditions consisted of acetonitrile and 20 mM ammonium formate in type 1 water pH 3 at a ratio of 92:8, respectively. Data acquisition and analysis were performed using Thermo Scientific Xcalibur 4.2.47. The analytical range was set from 10.0 to 50,000 ng/mL and the sample concentrations were interpolated from the standard curve.

Nirmatrelvir was extracted from mouse serum and tissue homogenates using protein precipitation as sample preparation technique. 180 µL of internal standard (IS) solution (100 ng/mL of EMD 281014 in methanol) was added to an aliquot of 20 µL of sample (serum or tissue homogenate). The sample was vortexed for approximately 5 seconds and let stand for a period of 10 minutes, then centrifuged at 16,000 x *g* for 10 minutes. One hundred and fifty microliters of the supernatant were transferred to a new tube and eight hundred and fifty microliters of 10 mM ammonium formate pH 3 buffer was added. The sample was vortexed for approximately 5 seconds and transferred to an injection vial for analysis. The analysis was performed using a Thermo Scientific TSQ Quantiva triple quadrupole mass spectrometer interfaced with a Thermo Scientific Ultimate 3000XRS UHPLC system using a heated electrospray ion source (HESI). MS detection was performed in positive ion mode, using selected reaction monitoring (SRM). The precursor-ion reactions for nirmatrelvir and IS were set to 500.3 → 319.1 and 377.2 → 209.1, respectively. A gradient mobile phase was used with a Thermo Scientific Accucore RP-MS analytical column (100 x 2.1 mm I.D., 2.6 µm) operating at 40°C. The initial mobile phase conditions consisted of acetonitrile and 10 mM ammonium formate in type 1 water pH 3.0 at a ratio of 10:90, respectively, and this ratio was maintained for 1 min. From 1 to 2.5 min a linear gradient was applied up to a ratio of 95:5 and maintained for 1.5 min. At 4.1 min, the mobile phase composition was reverted to the original conditions and the column was allowed to equilibrate for 5.9 min for a total run time of 10.0 min. Data acquisition and analysis were performed using ThermoScientific Xcalibur 4.2.47. The analytical range was set from 5.00 to 50,000 ng/mL and the sample concentrations were interpolated from the standard curve.

Molnupiravir in-life phase study was performed at Charles River Den Bosch, ‘s-Hertogenbosch, The Netherlands. Female BALB/c mice were dosed on a single occasion by oral gavage with molnupiravir formulated as a solution in 10% PEG400 and 2.5% Kolliphor-EL in Elix water. molnupiravir was administrated at doses of 3, 15 and 75 mg/kg to 3 groups of 6 animals. Blood samples (∼30 μL using K2EDTA-coated hematocrit capillaries) were collected from animals in all dose groups at 0.5, 1, 2, 6, 12 and 24 hours post dose by sampling from the jugular vein. Blood samples were centrifuged within 1 hour after blood sampling at approximately 3000g for 10 minutes at 2-8°C. Immediately after centrifugation, the plasma was transferred to a labeled polypropylene tube and stored in an ultra-low freezer set to maintain −80 ⁰C until shipment on dry ice to the bioanalytical laboratory. Plasma samples were then analyzed for molnupiravir metabolite EIDD-1931 with a standard HPLC-MS/MS method using an internal standard (Research Grade Assay 1).

#### Bioluminescence Imaging (BLI) of SARS-CoV-2 infection

All standard operating procedures and protocols for IVIS imaging of SARS-CoV-2-infected animals under ABSL-3 conditions were approved by IACUC, IBSCYU and YARC. All the imaging was carried out using IVIS Spectrum® (PerkinElmer) in XIC-3 animal isolation chamber (PerkinElmer) that provided biological isolation of anesthetized mice or individual organs during the imaging procedure. All mice were anesthetized via isoflurane inhalation (3 - 5 % isoflurane, oxygen flow rate of 1.5 L/min) prior and during BLI using the XGI-8 Gas Anesthesia System. Prior to imaging, 100 µL of nanoluc luciferase (nLuc) substrate, furimazine (NanoGlo^TM^, Promega, Madison, WI) diluted 1:40 in endotoxin-free PBS was retroorbitally administered to mice under anesthesia. The mice were then placed into XIC-3 animal isolation chamber (PerkinElmer) pre-saturated with isothesia and oxygen mix. The mice were imaged in both dorsal and ventral position at indicated days post infection. The animals were then imaged again after euthanasia and necropsy by spreading additional 200 µL of substrate on to exposed intact organs. Infected areas identified by carrying out whole-body imaging after necropsy were isolated, washed in PBS to remove residual blood and placed onto a clear plastic plate. Additional droplets of furimazine in PBS (1:40) were added to organs and soaked in substrate for 1-2 min before BLI.

Images were acquired and analyzed with Living Image v4.7.3 *in vivo* software package (Perkin Elmer Inc). Image acquisition exposures were set to auto, with imaging parameter preferences set in order of exposure time, binning, and f/stop, respectively. Images were acquired with luminescent f/stop of 2, photographic f/stop of 8. Binning was set to medium. Comparative images were compiled and batch-processed using the image browser with collective luminescent scales. Photon flux was measured as luminescent radiance (p/sec/cm2/sr). During luminescent threshold selection for image display, luminescent signals were regarded as background when minimum threshold setting resulted in displayed radiance above non-tissue-containing or known uninfected regions.

#### Plaque forming assay

Titers of virus stocks was determined by standard plaque assay. Briefly, the 4 x 10^5^ Vero-E6 cells were seeded on 12-well plate. 24 h later, the cells were infected with 200 µL of serially diluted virus stock followed by overlaying with 1ml of pre-warmed 0.6% Avicel (RC-581 FMC BioPolymer) made in complete RPMI medium. Plaques were resolved at 48 h post infection by fixing in 10 % paraformaldehyde for 15 min followed by staining for 1 hour with 0.2 % crystal violet made in 20 % ethanol. Plates were rinsed in water to visualize plaques.

#### Measurement of viral burden

Indicated organs (nasal cavity, brain, lungs) from infected or uninfected mice were collected, weighed, and homogenized in 1 mL of serum free RPMI media containing penicillin-streptomycin and homogenized in 2 mL tube containing 1.5 mm Zirconium beads with BeadBug 6 homogenizer (Benchmark Scientific, TEquipment Inc). Virus titers were measured using two highly correlative methods. Frist, the total RNA was extracted from homogenized tissues using RNeasy plus Mini kit (Qiagen Cat # 74136), reverse transcribed with iScript advanced cDNA kit (Bio-Rad Cat #1725036) followed by a SYBR Green Real-time PCR assay for determining copies of SARS-CoV-2 N gene RNA using primers SARS-CoV-2 N F: 5’-ATGCTGCAATCGTGCTACAA-3’ and SARS-CoV-2 N R: 5’-GACTGCCGCCTCTGCTC-3’. All the real-time PCR assays based on SYBR Green have a built-in melt-curve analyses to ensure estimation of only specific PCR products and not false-positives.

Second, we used nLuc activity as a shorter surrogate for plaque assay. Serially diluted clarified tissue homogenates were used to infect Vero-E6 cell culture monolayer. Infected cells were washed with PBS and then lysed using 1X Passive lysis buffer. The lysates transferred into a 96-well solid white plate (Costar Inc) and nLuc activity was measured using Tristar multiwell Luminometer (Berthold Technology, Bad Wildbad, Germany) for 2.5 seconds by adding 20 µl of Nano-Glo® substrate in nanoluc assay buffer (Promega Inc, WI, USA). Uninfected monolayer of Vero cells treated identically served as controls to determine basal luciferase activity to obtain normalized relative light units. The data were processed and plotted using GraphPad Prism 8 v8.4.3.

#### Analyses of signature inflammatory cytokines mRNA expression

Brain and lung samples were collected from mice at the time of necropsy. Approximately, 20 mg of tissue was suspended in 500 µL of RLT lysis buffer, and RNA was extracted using RNeasy plus Mini kit (Qiagen Cat # 74136), reverse transcribed with iScript advanced cDNA kit (Bio-Rad Cat #1725036). To determine mRNA copy numbers of signature inflammatory cytokines, multiplex qPCR was conducted using iQ Multiplex Powermix (Bio Rad Cat # 1725848) and PrimePCR Probe Assay mouse primers FAM-GAPDH, HEX-IL6, TEX615-CCL2, Cy5-CXCL10, Cy5.5-IFNgamma and HEX-IL1B. The reaction plate was analyzed using CFX96 touch real time PCR detection system. Scan mode was set to all channels. The PCR conditions were 95 °C 2 min, 40 cycles of 95 °C for 10 s and 60 °C for 45 s, followed by a melting curve analysis to ensure that each primer pair resulted in amplification of a single PCR product. mRNA copy numbers of *Il6, Ccl2, Cxcl10, Ifng* and *Il1b* in the cDNA samples of infected mice were normalized to *Gapdh* mRNA with the formula ΔC_t_(target gene)=C_t_(target gene)-C_t_(*Gapdh*). The fold increase was determined using 2^-ΔΔCt^ method comparing treated mice to uninfected controls.

#### Cryo-immunohistology of lung tissue

Lung tissues were isolated after necropsy and fixed in 1X PBS containing freshly prepared 4% PFA for 12 h at RT or 4 °C. They were then washed with PBS, cryoprotected with 10, 20 and 30% ascending sucrose series, snap-frozen in O.C.T.TM compound (Tissue-Tek) inside cryomolds and stored at −80C. 10 µm thick frozen sections were then cut and mounted on Superfrost plus slides. The sections were allowed to dry at 37 ° for 1 h and stored at −20 °C. For staining, the slides were equilibrated at RT, washed with 1X PBS to remove excess O.C.T and permeabilized with 0.2% Triton X-100 and treated with Fc receptor blocker (Innovex Biosciences) before staining with antibodies to mouse Krt8 followed by detection using secondary antibody conjugated to Alexa Fluor^TM^ 568 in PBS containing 2% BSA and 0.1% fetal bovine serum. Stained sections were treated with TrueVIEW Autofluorescence Quenching Kit (Vector Laboratories) and mounted in VECTASHIELD Vibrance Antifade Mounting Medium. Images were acquired and stitched using EVOS M7000 imaging system. The images were processed using Nikon Elements AR version 4.5 software (Nikon Instruments Inc, Americas) and figures assembled with Photoshop CC and Illustrator CC (Adobe Systems, San Jose, CA, USA).

#### Quantification and Statistical Analysis

Data were analyzed and plotted using GraphPad Prism software (La Jolla, CA, USA). Statistical significance for pairwise comparisons were derived by applying non-parametric Mann-Whitney test (two-tailed). To obtain statistical significance for survival curves, grouped data were compared by log-rank (Mantel-Cox) test. To obtain statistical significance for grouped data we employed 2-way ANOVA followed by Tukey’s multiple comparison tests. p values lower than 0.05 were considered statistically significant. P values were indicated as ∗, p < 0.05; ∗∗, p < 0.01; ∗∗∗, p < 0.001; ∗∗∗∗, p < 0.0001.

## Supplementary Figures

**Figure S1.**
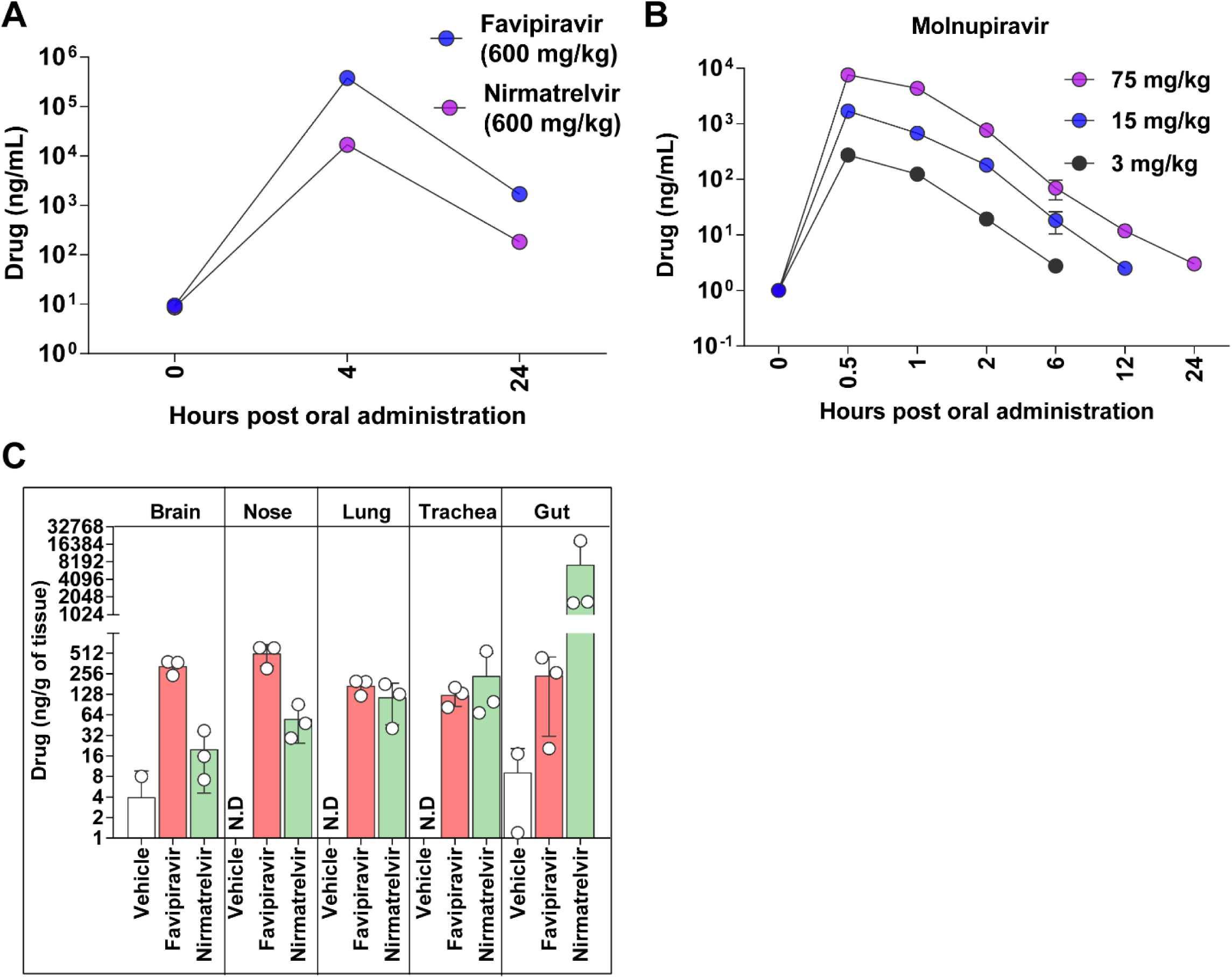
Pharmacokinetics of Antiviral Drugs used in the Study. Related to Figure 1. (A, B) Temporal concentrations (ng/ml) of indicated drugs in the serum. (A) favipiravir (600 mg/kg b.w, i.p.) and nirmatrelvir (600 mg/kg b.w., oral) were administered to K18-hACE2 mice and serum samples were collected at 0, 4 and 24 h post-treatment. (B) molnupiravir (3 mg/kg, 15 mg/kg and 75 mg/kg b.w, oral) was administered to BALB/c mice and serum samples were collected at 0.5, 1, 2, 6, 12 and 24 h post-treatment. (C) Drug concentrations (ng/g of tissue) in indicated clarified tissue homogenates for an experiment as in A at 24 h post-treatment following necropsy.

**Figure S2.**
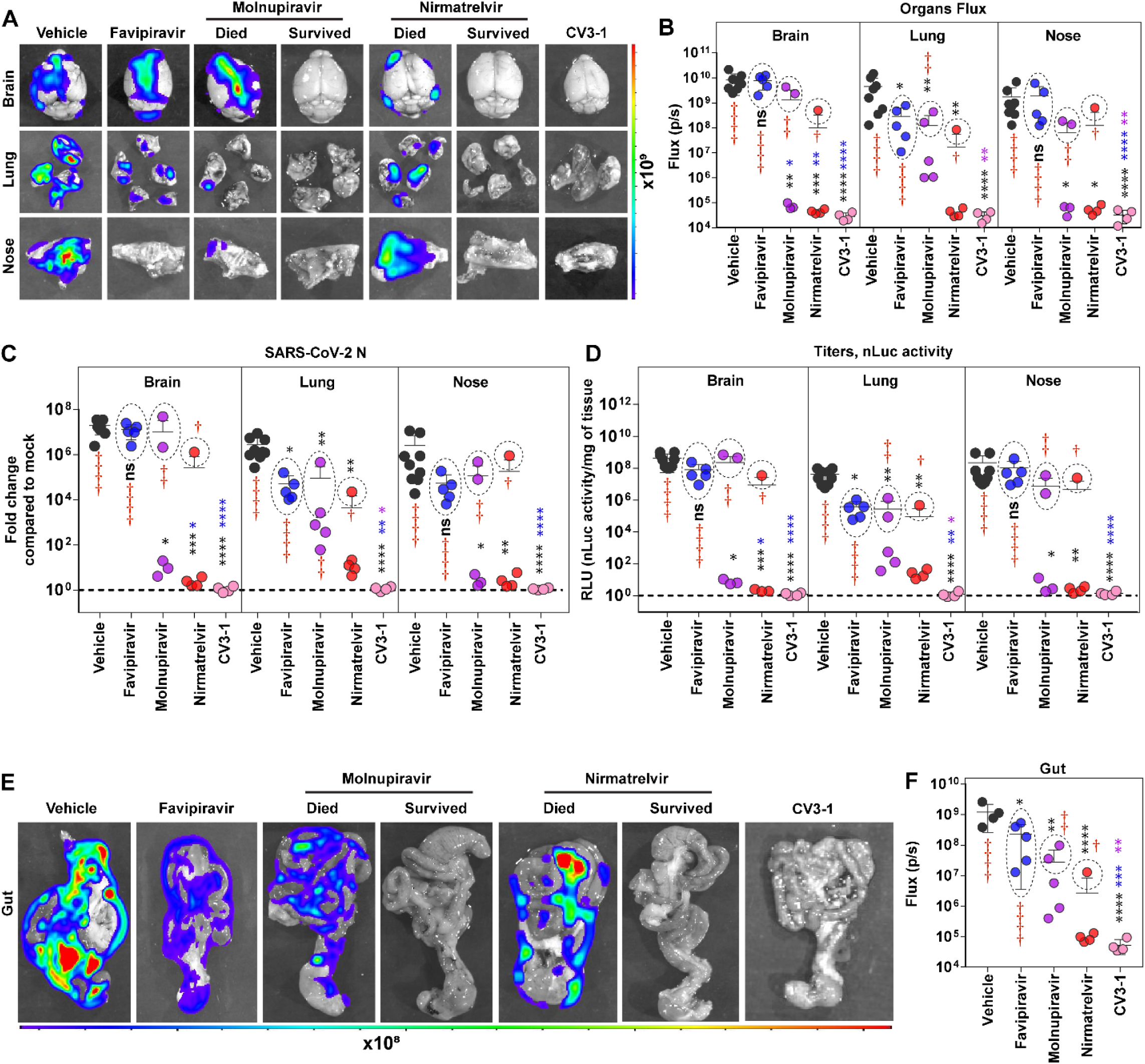
In Vivo Efficacy of Antiviral Drugs and Neutralizing Antibody Against Delta VOC Related to Figure 1. (A-B) *Ex vivo* imaging of indicated organs and quantification of nLuc signal as flux (photons/sec) after necropsy for an experiment shown in Figure 1A. Organs from dead and surviving mice are shown separately for clarity. (C) Fold changes in nucleocapsid (N) mRNA expression in brain, lung, and nasal cavity. Data were normalized to *Gapdh* mRNA expression in the same sample and that in non-infected mice after necropsy when mice died from infection or at 14 or 22 dpi for surviving mice. (D) Viral loads (nLuc activity/mg) in indicated organs from mice under specified treatment regimens evaluated using Vero E6 cells as targets when mice died from infection or at 14 or 22 dpi for surviving mice. (E, F) *Ex vivo* imaging of gut and quantification of nLuc signal as flux (photons/sec) after necropsy at the time of death (see Figure 1F) for an experiment shown in Figure 1A. Grouped data in (B-D), were analyzed by 2-way ANOVA followed by Tukey’s multiple comparison tests. The data in (F) was analyzed by one-way ANOVA followed by Kruskal-Wallis’ test. Statistical significance for group comparisons to Vehicle are shown in black, with favipiravir are shown in blue, with molnupiravir are shown in purple, with nirmatrelvir are shown as red and with CV3-1 are shown as pink. ∗, p < 0.05; ∗∗, p < 0.01; ns not significant; Mean values ± SD are depicted. Dotted circles with red cross signs at indicated time points are used to demarcate mice that succumbed to infection.

**Figure S3.**
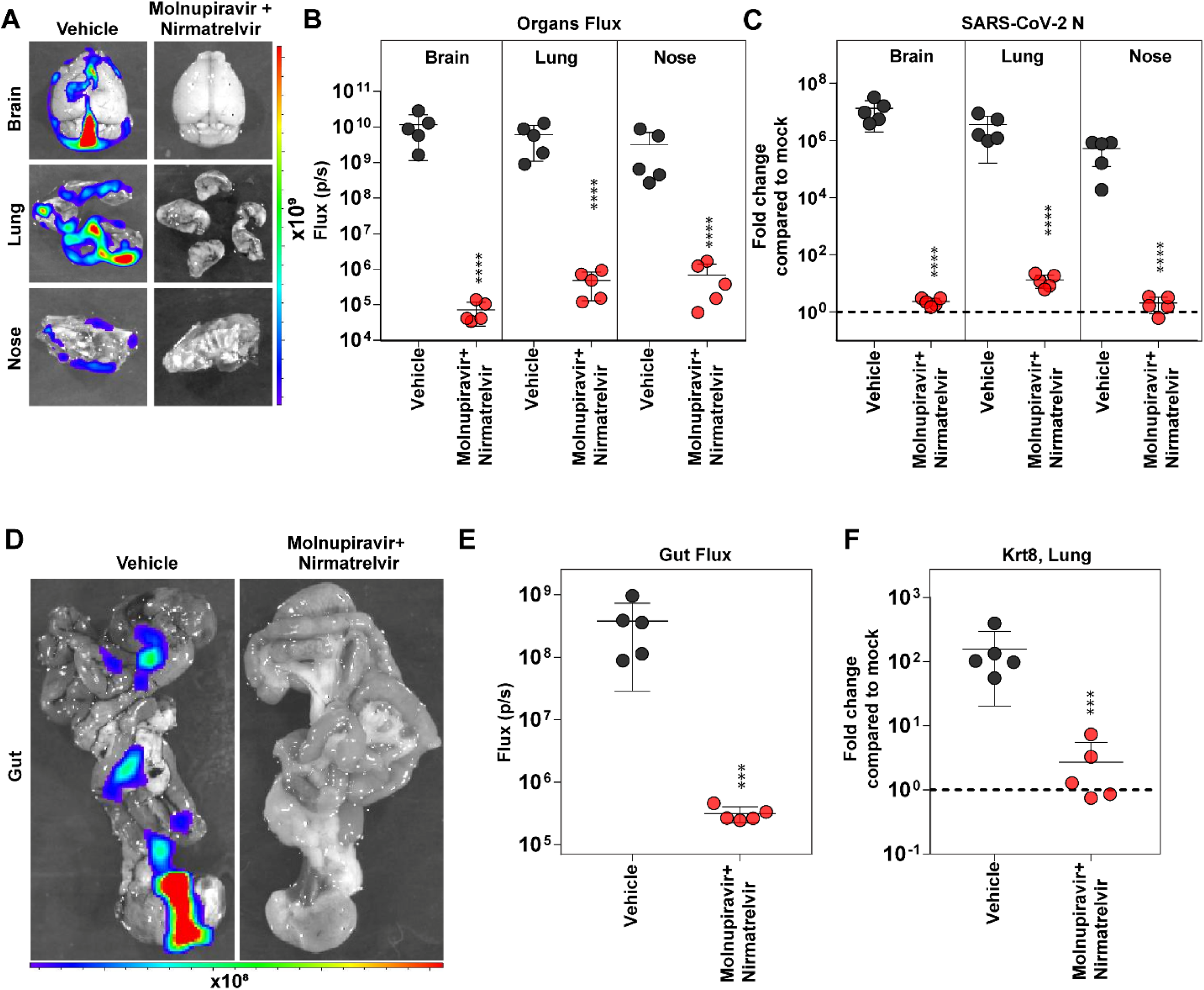
*In Vivo* Efficacy of Molnupiravir Combined with Nirmatrelvir Against Delta VOC. Related to Figure 2. (A-B) *Ex vivo* imaging of indicated organs and quantification of nLuc signal as flux (photons/sec) after necropsy for an experiment shown in Figure 2A. (C) Fold changes in nucleocapsid (N) mRNA expression in brain, lung, and nasal cavity upon death or at 14 dpi in the surviving animals. Data were normalized to *Gapdh* mRNA in the same sample and that in non-infected mice after necropsy. (D, E) *Ex vivo* imaging of gut and quantification of nLuc signal as flux (photons/sec) after necropsy at the time of death (see Figure 2F) for an experiment shown in Figure 2A. Grouped data in (B-D), were analyzed by 2-way ANOVA followed by Tukey’s multiple comparison tests. The data in (F) was analyzed by no parametric Mann-Whitney test. ∗, p < 0.05; ∗∗, p < 0.01; ns not significant; Mean values ± SD are depicted.

**Figure S4.**
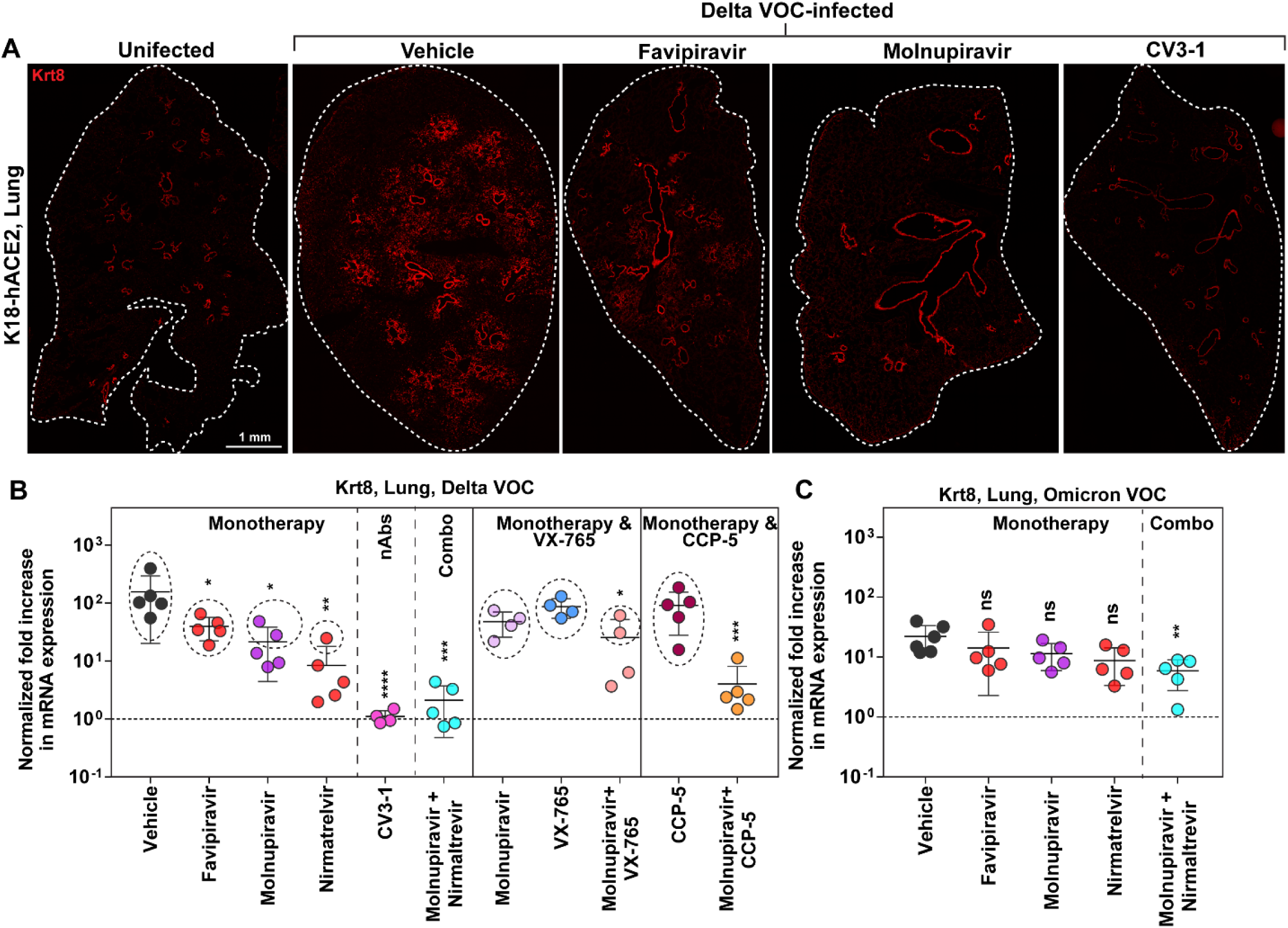
Therapeutic Interventions using Drug Monotherapy or Evaluated Combinations Reduced SARS-CoV-2 VOC-mediated Krt8 Expression, a Marker for Lung Injury. Related to Figures 1-6. (A) Images of lung cryosections from uninfected or Delta VOC-challenged K18-hCE2 mice that were treated as indicated and immunostained with antibodies to mouse Krt8 followed by detection using secondary antibody conjugated to Alexa Fluor^TM^ 568 for an experiment as in Figure 1A. Enhanced Krt8 staining of injured alveolar epithelial cells due to Delta VOC infection in the areas surrounding the bronchiolar epithelium can be clearly seen in Vehicle and favipiravir-treated mouse lung compared to predominant staining of bronchial epithelial cells in CV3-1 nAb and molnupiravir (survived) treated conditions. Size bar: 1 mm (B, C) Summary of fold changes in cytokeratin 8 (*Krt8*) mRNA expression in the lung of mice infected with Delta or Omicron nLuc reporter VOCs under specified therapeutic interventions at the time of death or 14-22 dpi for surviving animals described in the study. Data were normalized to *Gapdh* mRNA expression in the same sample and that in non-infected mice after necropsy. Each data point represents one mouse. Dotted circles are used to denote mice that succumbed to infection. Grouped data in (A-B), were analyzed by 2-way ANOVA followed by Tukey’s multiple comparison tests for determining statistical significance to vehicle-treated mice. ∗, p < 0.05; ∗∗, p < 0.01; ns not significant; Mean values ± SD are depicted.

**Figure S5.**
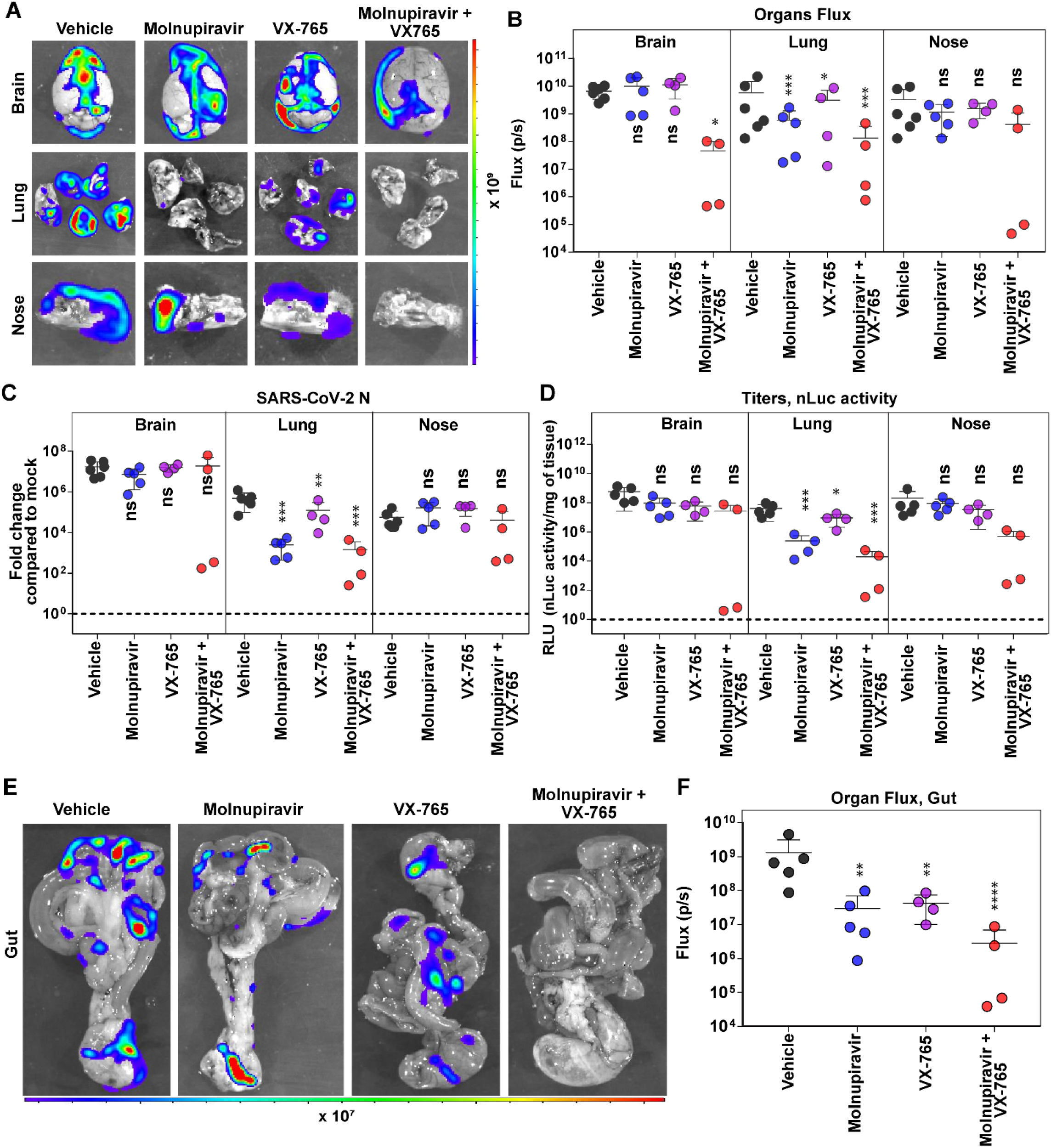
*In Vivo* Efficacy of Molnupiravir Combined with VX-765 Against Delta VOC Related to Figure 5. (A-B) *Ex vivo* imaging of organs from mice under indicated treatment regimens and quantification of nLuc signal as flux (photons/sec) after necropsy for an experiment shown in Figure 5A. (C) Fold changes in nucleocapsid (N) mRNA expression in brain, lung, and nasal cavity tissues upon death or at 14 dpi in the surviving animals. Data were normalized to *Gapdh* mRNA in the same sample and that in non-infected mice after necropsy. (D) Viral loads (nLuc activity/mg) in tissues from mice under specified treatment regimens using Vero E6 cells as targets upon death or at 14 dpi for surviving mice. (E, F) *Ex vivo* imaging of gut and quantification of nLuc signal as flux (photons/sec) after necropsy at the time of death (see Figure 5F) or at 14 dpi for surviving mice for an experiment shown in Figure 5A. Grouped data in (B-D), were analyzed by 2-way ANOVA followed by Tukey’s multiple comparison tests. The data in (F) was analyzed by one-way ANOVA followed by Kruskal-Wallis’ test. Statistical significance for group comparisons to Vehicle are shown in black, with molnupiravir are shown in blue, with VX-765 are shown in purple and with molnupiravir combined with VX-765 are shown as red. ∗, p < 0.05; ∗∗, p < 0.01; ∗∗∗, p < 0.001; ∗∗∗∗, p < 0.0001; ns not significant; Mean values ± SD are depicted.

**Figure S6.**
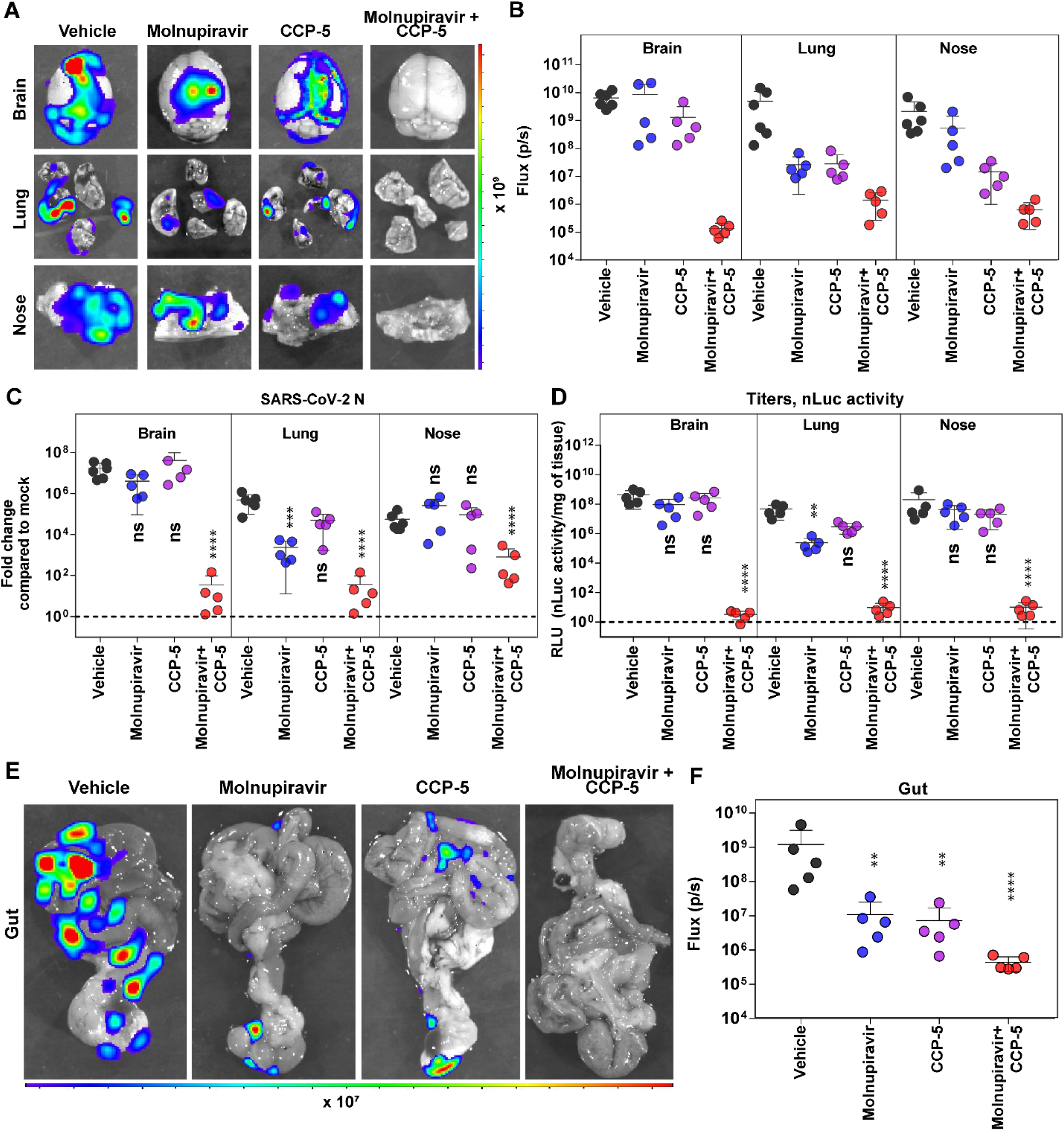
In Vivo Efficacy of Molnupiravir Combined with CCP-5 Against Delta VOC Related to Figure 6. (A-B) *Ex vivo* imaging of organs from mice under indicated treatment regimens and quantification of nLuc signal as flux (photons/sec) after necropsy for an experiment shown in Figure 6A. (C) Fold changes in nucleocapsid (N) mRNA expression in brain, lung, and nasal cavity upon death or at 14 dpi in the surviving animals. Data were normalized to *Gapdh* mRNA in the same sample and that in non-infected mice after necropsy. (D) Viral loads (nLuc activity/mg) in tissues from mice under specified treatment conditions using Vero E6 cells as targets upon death or at 14 dpi for surviving mice. (E, F) *Ex vivo* imaging of gut and quantification of nLuc signal as flux (photons/sec) after necropsy at the time of death or at 14 dpi for surviving mice (see Figure 6F) for an experiment shown in Figure 6A. Grouped data in (B-D), were analyzed by 2-way ANOVA followed by Tukey’s multiple comparison tests. The data in (F) was analyzed by one-way ANOVA followed by Kruskal-Wallis’ test. Statistical significance for group comparisons to Vehicle are shown in black, with molnupiravir are shown in blue, with CCP-5 are shown in purple and with molnupiravir with CCP-5 are shown as red. ∗, p < 0.05; ∗∗, p < 0.01; ∗∗∗, p < 0.001; ∗∗∗∗, p < 0.0001; ns not significant; Mean values ± SD are depicted.

## STAR Methods

### KEY RESOURCES TABLE

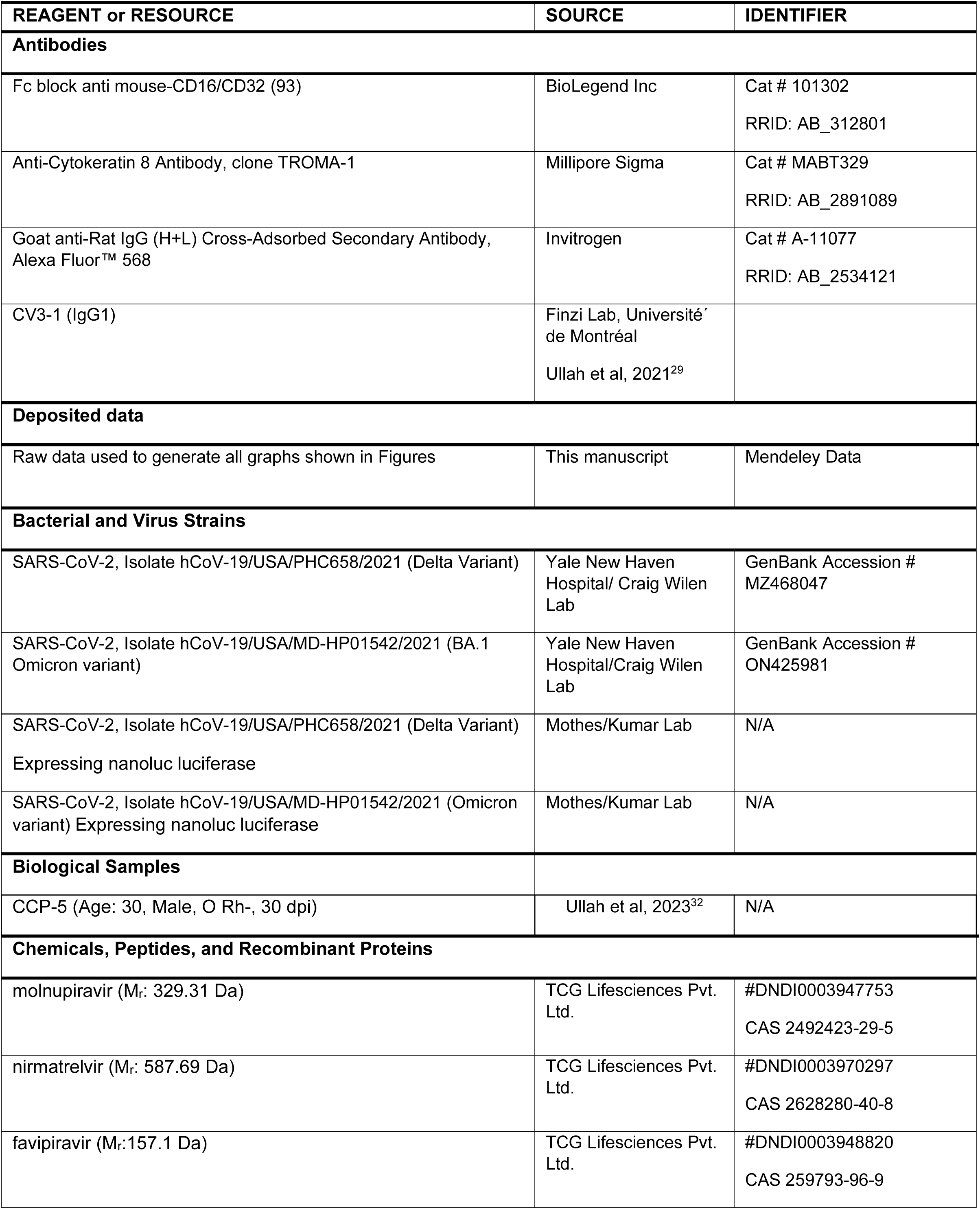
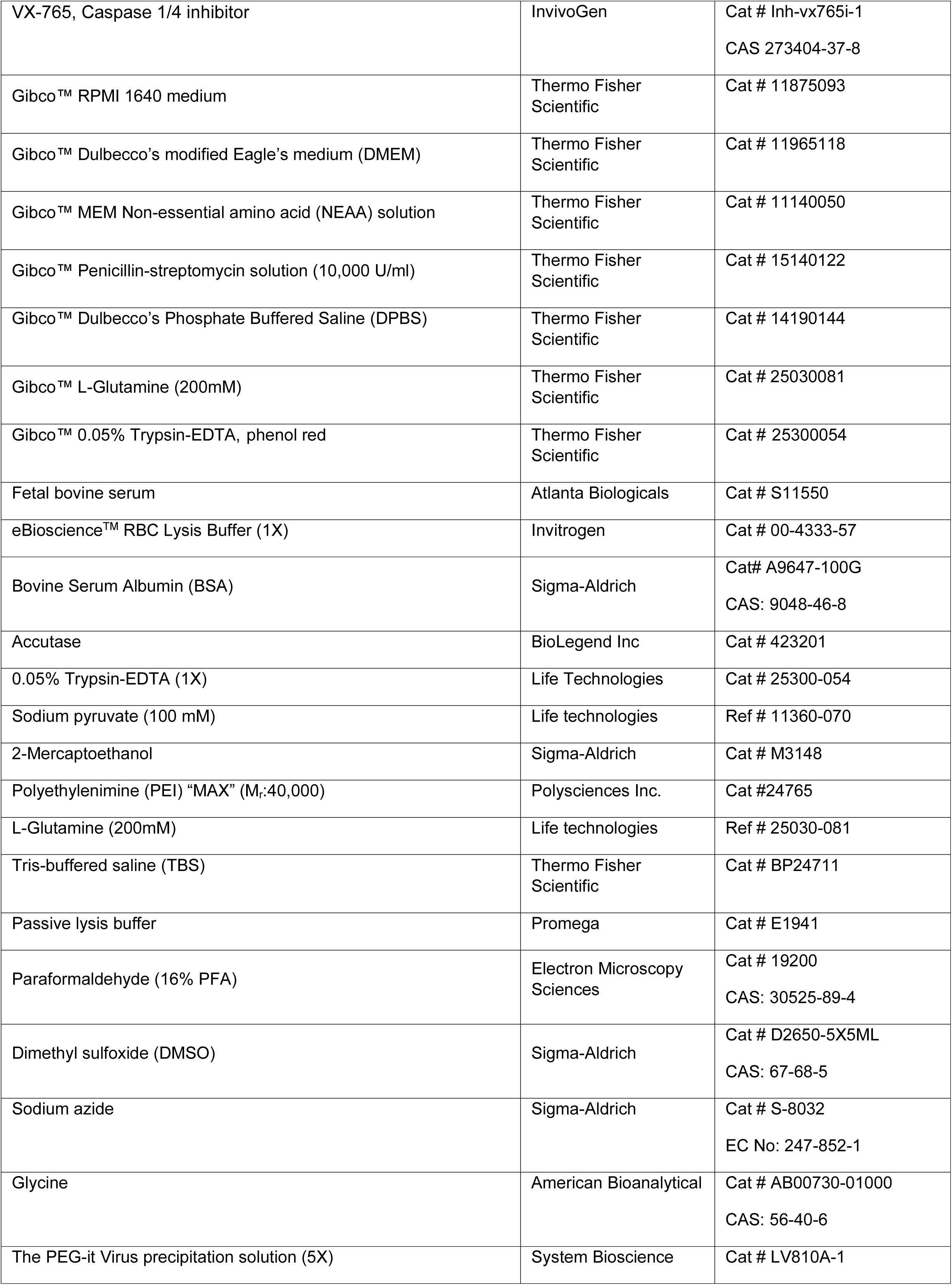
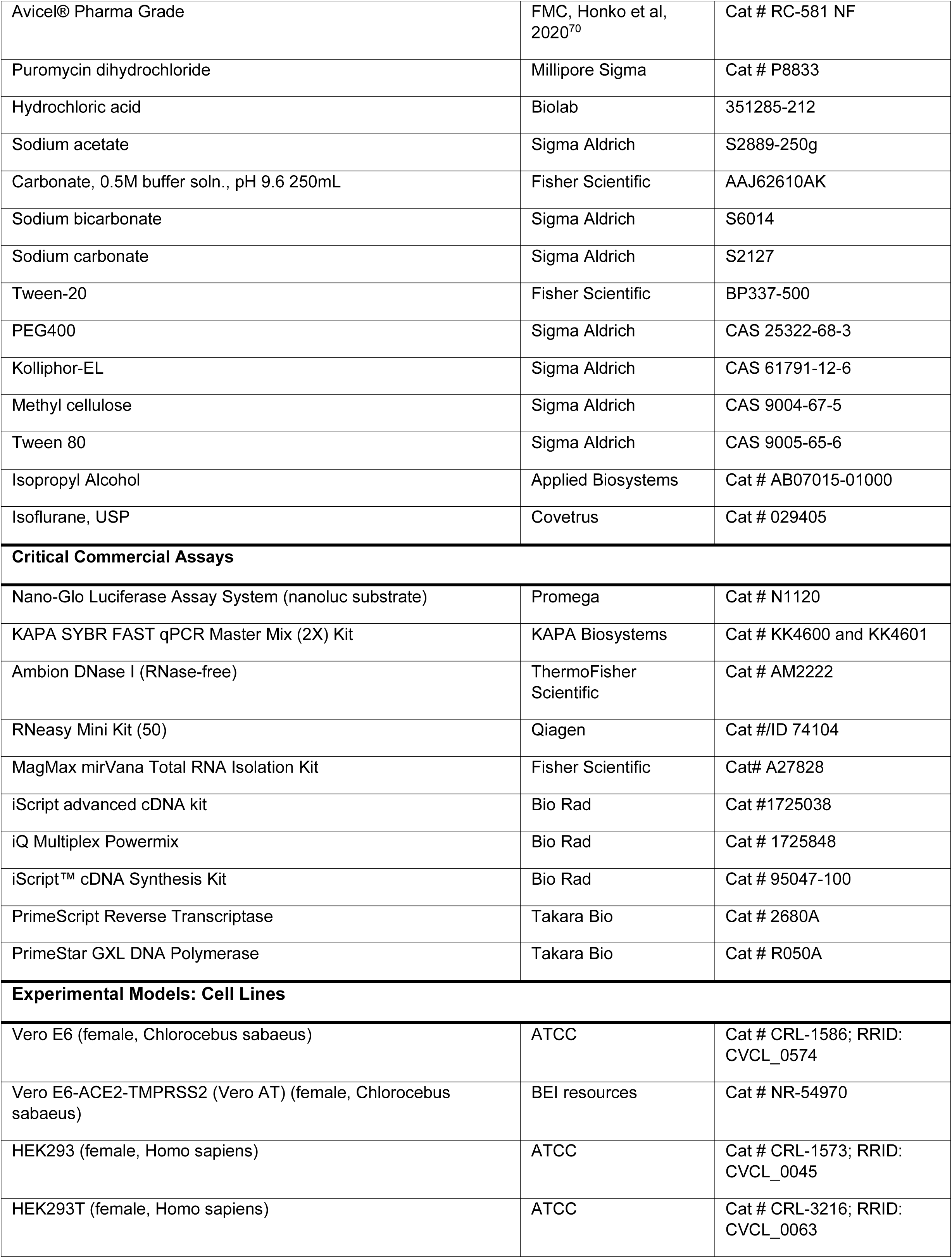
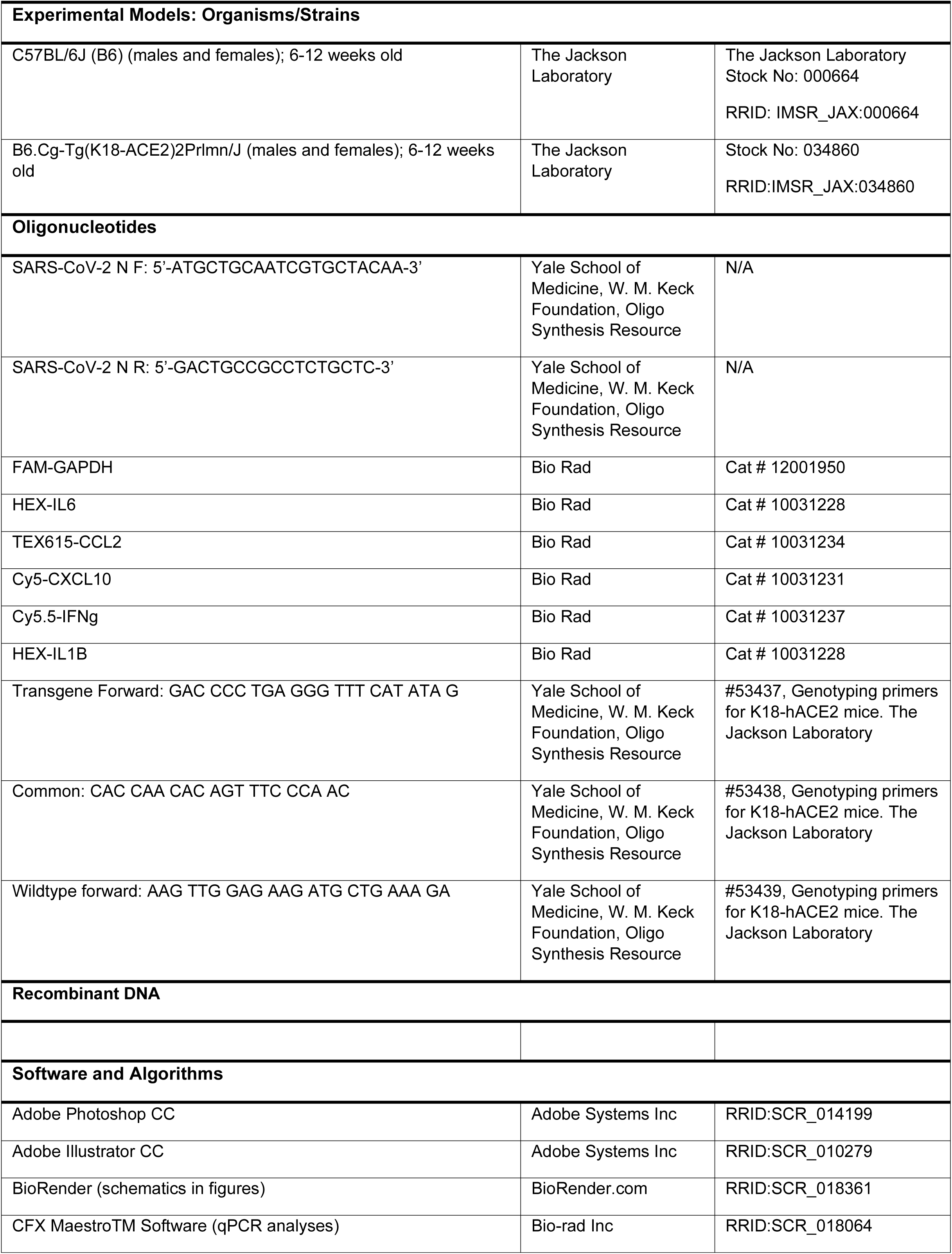
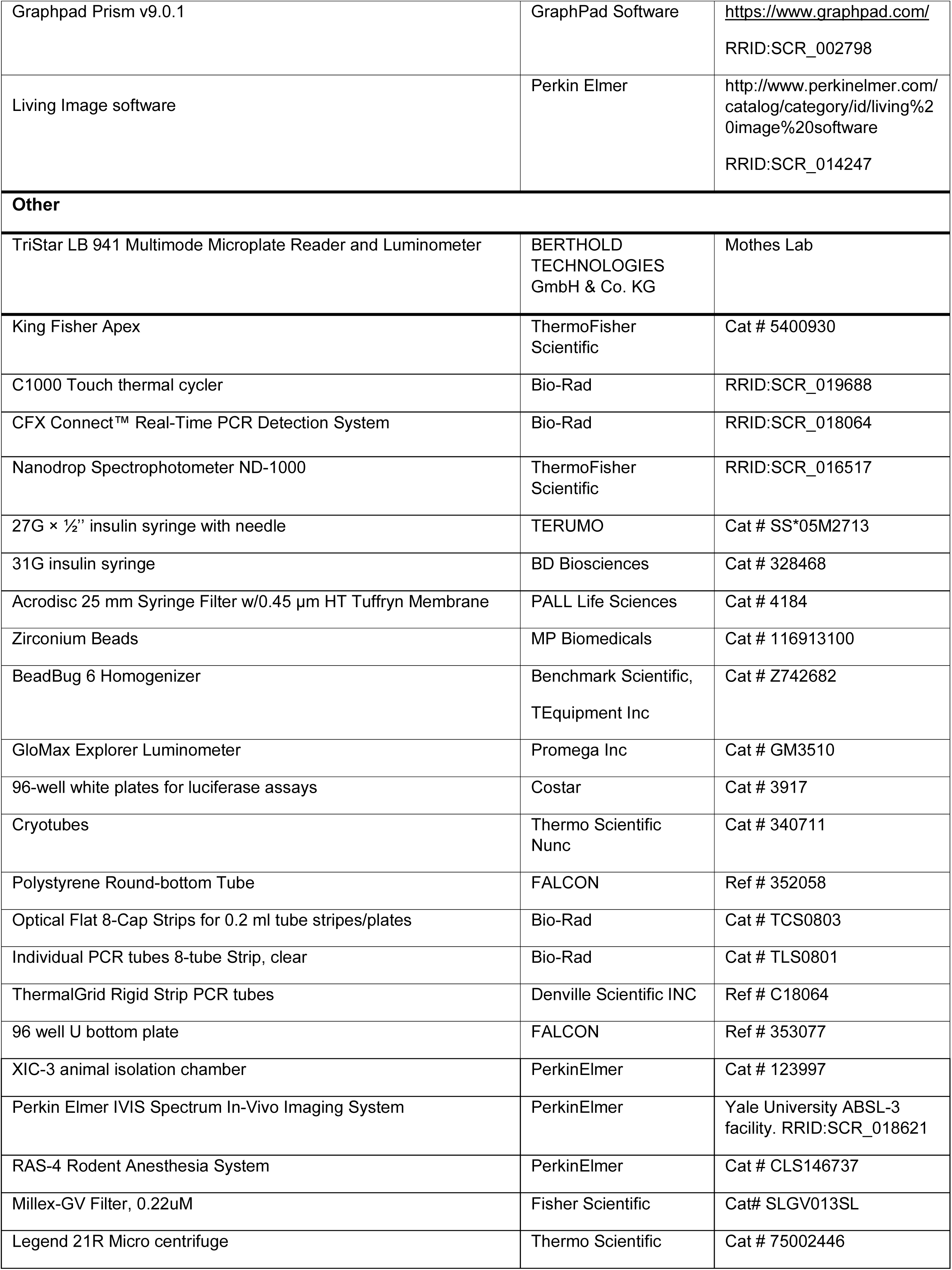
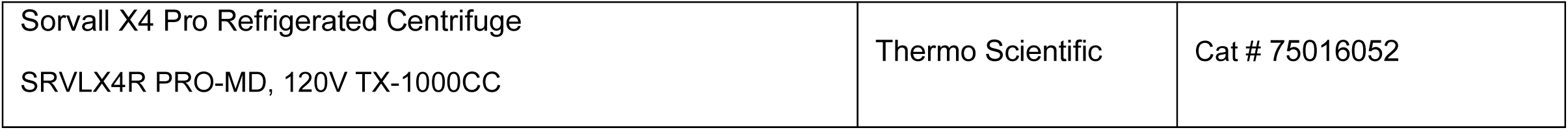

## Notes

### Competing Interest Statement

The authors have declared no competing interest.

